# Pericytes mediate neurovascular remodeling in chronic arterial hypertension

**DOI:** 10.1101/2024.05.13.594041

**Authors:** Lorena Morton, Alejandra P. Garza, Grazyna Debska-Vielhaber, Luis E. Villafuerte, Solveig Henneicke, Philipp Arndt, Sven G. Meuth, Stefanie Schreiber, Ildiko R. Dunay

## Abstract

Chronic arterial hypertension restructures the vascular architecture of the brain, leading to a series of pathological responses that culminate in cerebral small vessel disease. Pericytes respond dynamically to vascular challenges; however, how they manifest under the continuous strain of hypertension has not been elucidated. Therefore, in this study, we characterized pericyte behavior alongside hypertensive states in the spontaneously hypertensive stroke-prone rat (SHRSP) model, emphasizing their phenotypic and metabolic transformation. Our results reveal an early transition in PDGFRß^+^ pericytes toward increased NG2 and CD13 co-expressing subtypes, signaling enhanced pericyte reactivity in an effort to stabilize vascular structures and an inflammatory engagement within the vascular niche in response to hypertensive stress. Gene expression profiling of microvessels revealed altered expression within crucial pathways i.e., angiogenesis, blood-brain barrier integrity, hypoxia and inflammation. Furthermore, we detected that circulating extracellular vesicles from SHRSP alter pericyte mitochondrial membrane potential, highlighting their ability to transmit pathogenic signals that exacerbate vascular remodeling. Detailed metabolic analysis revealed a significant shift toward glycolytic metabolism in pericytes already in initial hypertension, alongside a dysregulation of ATP production pathways. These findings emphasize the transformative influence of hypertension on cerebral pericytes and the extensive consequences on cerebral vascular health.

## Introduction

Cerebral microvasculature is essential for brain health, playing a pivotal role in maintaining vascular stability and homeostasis^1–6^. Pericytes, central to the integrity of the blood-brain barrier (BBB)^7, 8^, regulate blood flow^9,10^, angiogenesis^11^, and orchestrate vascular immune homeostasis of the neurovascular unit (NVU)^12–15^, thereby collectively contributing to immunosurveillance and overall cerebrovascular health. Their dysfunction has been implicated in a range of cerebrovascular diseases, from stroke to dementia, indicating their importance across various pathologies^16–21^. In conditions such as chronic arterial hypertension^22^, pericyte dysfunction may be linked to BBB disruption by endothelial cell malfunction, cerebrovascular remodeling, and the onset of cerebral small vessel disease (cSVD)^23, 24^. cSVD is characterized by white matter lesions, vascular-associated activated microglia states, and microvascular abnormalities, exacerbated by chronic inflammation associated with aging and chronic arterial hypertension^25–29^. Among its subtypes, arteriolosclerosis, marked by thickening and stiffening of vessel walls, is prevalent in the elderly and in those with vascular risk factors, significantly altering cerebral microcirculation and heightening ischemic vulnerability^30^.

Recent focus on extracellular vesicles (EVs) have gained attention for their role in cellular communication, influencing the behavior of cells through the transfer of bioactive molecules^31, 32^. These vesicles, which are shed by all cell types into the circulation, facilitate a unique cellular interaction by carrying a cargo of proteins, lipids, and nucleic acids, capable of modulating recipient cell function and contributing to disease processes^33^.

Pericytes respond dynamically to vascular challenges; however, the specific responses under the continuous strain of hypertension has not been elucidated. Here, using the spontaneously hypertensive stroke-prone rat (SHRSP)^34, 35^, which closely mimics hypertensive cSVD in humans^24, 36–38^, we examine how pericytes adapt to prolonged arterial hypertension. We demonstrate how pericytes respond to hypertensive stress by altering their phenotype and metabolic pathways early in disease onset. We revealed that in the face of systemic hypertension, pericytes increase their dependence on glycolytic pathways, a metabolic shift that might initially serve adaptive purposes but becomes maladaptive. This shift is accompanied by significant changes in the expression of platelet-derived growth factor receptor-ß (PDGFRß), neuroglial antigen 2 (NG2), and ANPEP (CD13), canonical pericyte markers^39, 40^. We also provide novel evidence on the metabolic reprogramming of pericytes in chronic arterial hypertension and the involvement of EVs in mediating vascular pathology across the NVU.

## Materials and Methods

### Animals

All experiments adhered to the German Animal Welfare Ordinance and were approved by the Animal Care Committee of Saxony-Anhalt (license identification numbers 42502-2-1277 for Uni MD). A total of 116 male rats were used across various experimental phases, as detailed in Suppl. Data Table 1. For the initial hypertension (HTN) phase, we refer to 6-8 weeks old Wistar (Ctrl) and spontaneously hypertensive stroke-prone rats (SHRSP). The early chronic hypertension phase includes 25-weeks old rats, and the late chronic hypertension phase comprises 34-36 week old rats. All animals were sourced from Charles River Laboratories (Wilmington, Massachusetts, United States for SHRSP; Research Models and Services, Germany GmbH, Sulzfeld, Germany for Wistar). Wistar rats served as controls, while SHRSP rats model hypertension at specified ages, which reflect the progression from initial to late chronic hypertension ^35, 36^. All animals were housed with a natural light-night cycle, had access to water and food *ad libitum* and were monitored daily to assess neurological functions. SHRSP develop a vascular risk profile characterized by arterial hypertension from 6 weeks of age and develop classical cSVD pathology^38, 41, 42^.

### Vascular cell isolation for flow cytometry

To characterize pericytes and endothelial cells, vascular cells were isolated from 6-, 25-, and 34-week-old Ctrl and HTN rats using a modified version of Crouch et al., adapted for rat brains^43^. Following anesthesia with pentobarbital and hemisphere preparation, cerebral cortices were enzymatically dissociated with collagenase/dispase, mechanically triturated, and purified using a 22% Percoll gradient (Cat. no. GE17-0891-01, Sigma-Aldrich). Isolated cells were washed, counted, centrifuged, resuspended, and stained. Unstained cells, Fluorescence Minus One (FMO), and single-staining controls with compensation beads were used as controls. Prior to fluorochrome-conjugated antibody labeling, cells were incubated with purified mouse anti-rat CD32 FcγII (clone D34-485, Rat BD Fc Block) for unspecific binding. Cell viability was assessed using a dead dye (Zombie NIR™, Biolegend), and cells were stained with a panel of fluorophore-conjugated antibodies, including anti-CD45 (clone: OX-1), anti-CD31 (clone: TLD-3A12), anti-Pdgfrß (clone: rabbit polyclonal), anti-NG2 (clone: D120.43), and anti-CD13 (clone: WM15), followed by fixation, permeabilization, and anti-Ki-67 (clone: SolA15) staining. Data were acquired and analyzed using AttuneNxT (Thermo Fisher Scientific) and Flowjo Analysis Software v10.5.3 (BD). Uniform Manifold Approximation and Projection (UMAP) analysis was performed on exported live, single, CD45-negative, CD31, and PDGFRß-positive cells downsampled to 7,500 cells per group and time point. Concatenated data were projected to UMAP using vascular compensated markers and 25 nearest neighbors with a minimum distance of 0.8 as running parameters.

### Tissue dissociation and microvessels isolation

Brains from Ctrl and HTN rats aged 8, 25, and 34 weeks were harvested following transcardial perfusion with PBS-EDTA for microvessels isolation utilizing already established optimized protocols^29, 44, 45^. Briefly, cortices were mechanically dissociated by mincing the tissue into fine fragments and enzymatically digested with Collagenase Type II (1 mg/ml) and DNase I (15 µg/ml) in DMEM/F12 containing penicillin (100 units/ml), streptomycin (100 µg/ml), and glutamine (2mM). Myelin and neurons were removed via centrifugation in 20% BSA-DMEM/F12 at 1000 g for 20 min, repeating the process three times. Microvessels were collected from pellets into new sterile tubes and further digested with collagenase-dispase (1 mg/ml) and DNase I (6.7 μg/ml) in DMEM for 60 min at 37 °C. Microvessels were layered on top of a 33% Percoll (Cat. no. GE17-0891-01, Sigma-Aldrich) density gradient and centrifuged for 10 min at 1000 g at 4 °C. Microvessels were washed twice, filtered through a 40 µm cell strainer, reversed, washed into a new sterile tube, and centrifuged at 400 g for 10 min at 4 °C. Microvessels were assessed microscopically before being stored in RNAlater (AM7020, Thermo Fisher Scientific) for further processing, or seeded for *in vitro* culture assays for pericyte *in vitro* expansion.

### RNA isolation and RT^2^ profiler PCR array

Collected microvessels were pelleted at 20,000 × g for 10 min and resuspended in 350 μl of RLT Plus Buffer from the RNeasy Micro Kit (QIAGEN). RNA was isolated using the RNeasy Plus Micro Kit, following the manufacturer’s instructions, as previously described^29^. RNA quality was determined using a NanoDrop 2000 (Thermo Fisher Scientific) spectrophotometer and used in combination with Power SYBR® Green RNA-to-CT™ 1-Step Kit (Cat. no. 4389986, Thermo Fisher Scientific) on a custom RT^2^ profiler PCR Array (QIAGEN) using a LightCycler® 96 (Roche). Cycle Threshold (C_T_) values were exported to an Excel file to create a table. This table was then uploaded to the data analysis web portal at http://www.qiagen.com/geneglobe. Samples were assigned to control and test groups. C_T_ values were normalized based on the selection of reference genes. The data analysis web portal calculated fold change and gene regulation using the 2−ΔΔCt method^46^. *Hprt* gene was used to normalize the results. Data were further normalized to the respective mean level in age-matched early Ctrl. The full list of genes included in the custom PCR array is provided in Suppl. Data Table 2. Downstream GO analyses, data presentation, and statistical analyses of exported gene regulation tables were performed using STRING database version 12.0 and GraphPad Prism 10.

### Pericyte *in vitro* subculture

Rat cerebral pericytes were cultured from isolated brain microvessels containing a mix of pericytes and endothelial cells. Initial microvessel seeding was established in DMEM/F12 supplemented with 15% platelet-derived serum, 1 ng/ml bFGF, and 100 µg/ml heparin for the first 2 days. The medium was then switched to DMEM supplemented with 10% FBS, 1% non-essential amino acids, and 1% penicillin/streptomycin to support pericyte survival and proliferation. To enrich pericyte populations and eliminate endothelial cells, cultures were treated in the absence of puromycin, allowing pericyte expansion. Subsequent cell expansions were done using Accutase® Cell Solution (Biolegend) for gentle detachment, ensuring optimal cell viability. For cryopreservation, cultures were frozen in CryoStor® CS10 (Stemcell Technologies) and stored in liquid nitrogen.

### Immunofluorescence staining

Pericytes were expanded *in vitro* from seeded 8-week-old rat microvessels. At passages 11-13 pericytes at 90-95% confluence from T-75 cell culture flasks (Greiner CELLSTAR), were seeded at a density of 20,000 cells/well in 12-well plates containing 20 mm coverslips. Coverslips were sterilized and coated with rat tail collagen 24 h prior to pericyte expansion. Cells were fixed with 4% PFA and washed 48 h after seeding. Thereafter, cells were blocked with 3% horse serum for 2 h. Primary antibodies were incubated overnight at room temperature in the same solution. After PBS washes, cells were incubated with secondary antibodies diluted 1:1000 in 0.1% Triton X-100. Mounting was performed with ProLong mounting medium containing DAPI (Cat. no. P36935, Thermo Fisher Scientific), and coverslips were dried in darkness before microscopy analysis. Imaging was performed to visualize and confirm pericyte identity and purity using a TCS SP8 X Laser Confocal Microscope (Leica Microsystems) software (LAS-AF 1.8.1.13759). Full list of antibodies and their specificities can be found in Suppl. Data Table 3.

### Immunofluorescence image processing and analysis

Immunofluorescence images were processed using ImageJ (Rasband, W.S., (https://imagej.nih.gov/ij/, 1997-2018)), and viewed as hyperstacks in default color mode. The scale was changed from grayscale to specific colors for each channel. Channels were split, with contrast and brightness adjusted uniformly across samples to maintain consistent parameters. Individual adjusted and composite images created were saved as TIFF files. For z-stack images, after color adjustment and maximum intensity projection, images were processed similarly by splitting and merging channels, as described above, with the final composite image converted to RGB and saved as TIFF. The Mean Fluorescence Intensity (MFI) of PDGFRβ, NG2, and CD13 from individual cells was exported from processed images, as described previously^47^. Quantitative data were used to plot the MFI for each marker, allowing for the analysis of expression patterns in control versus hypertensive pericytes.

### Flow cytometric analysis of *in vitro* pericytes

At passages 11-13 pericytes at 90-95% confluence from T-75 cell culture flasks were dissociated with Accutase® Cell Solution (BioLegend) and prepared for flow cytometry to analyze subtype distribution *in vitro*, comparing control vs. hypertensive-derived cells. Cells were stained with antibodies against PDGFRβ, NG2, and CD13 and analyzed using an Attune NxT Flow Cytometer. FMO controls and unstained samples were used to set gates. Post-acquisition data analysis was performed using FlowJo software (v10.5.3). Data were biexponentially transformed for scaling and t-distributed Stochastic Neighbor Embedding (tSNE) was employed on compensated PDGFRβ-positive cells at a perplexity of 10 and 1,500 iterations to examine the differential expression of CD13 and NG2 in control versus hypertensive pericytes.

### Plasma-derived extracellular vesicles

Peripheral blood was collected from the portal vein of anesthetized animals using a 21G butterfly needle, immediately mixed with acid citrate dextrose (ACD), and subjected to two inversions for agent incorporation. Samples were processed within 1 h, as previously described^48^. Briefly, plasma separation was initially achieved by centrifugation at 1500 × g for 10 min, which was then transferred to sterile 1.5 ml Eppendorf tubes. To isolate EVs, the plasma underwent two centrifugation cycles at 1500 × g for 10 min maintaining supernatant, followed by two rounds of ultracentrifugation at 14,000 × g for 70 min, discarding supernatants and resuspending pellets in 0.22 µm filtered PBS (fPBS -/-). The final EV pellet was resuspended in 200 µL of fPBS -/- and stored at −80 °C. For EV characterization, 10 µL of isolated EVs was then mixed with 90 µL of fPBS -/-, and samples were labeled with CD9, CD63, and CD81 (APC-conjugated, Clones HI9A, H5C6 and 5A6, respectively, Biolegend). Samples were analyzed using an Attune NxT Flow Cytometer equipped with a small particle side scatter filter. Size gating (300-1000 nm) was established using silica beads (Creative Diagnostics). Acquisition of samples was performed at a speed of 25 µL per min, with SSC threshold set to 0.18 x 10^3^ and FSC threshold set to 0.15 x 10^3^. A stop option was activated upon reaching 150,000 events within the size gate. Each sample measurement was followed by a 10% bleach and 0.22 µm filtered distilled water rinse. Data analysis was conducted using FlowJo 10.9.0 and GraphPad Prism 10 for subsequent downstream analyses.

### JC-10 Mitochondrial membrane potential assay

Enriched pericytes at 90-95% confluence from T-75 cell flasks derived from control rats were expanded and seeded in 96-well plates at a density of 10,000 cells/well for 24 h prior. Cells were subjected to a JC-10 mitochondrial membrane potential assay (Cat. no. 421902, BioLegend) to assess metabolic responses to plasma and plasma-derived extracellular vesicles (pdEVs). Before stimulation cells were incubated in pericyte ultracentrifuged (18 h × 110,000 g) basal medium containing 1% FCS. Treatments included 10% v/v normotensive or hypertensive plasma, or normotensive or hypertensive pdEVs standardized to a protein concentration of 13.5 µg. All stimulations were induced for 6 h. JC-10 dye was added following the manufacturer’s instructions, and changes in mitochondrial membrane potential were measured by calculating the ratio of red/green fluorescence ratio indicative of mitochondrial health using the AttuneNxT and Flowjo Analysis Software (v10.5.3).

### Extracellular flux analysis

Metabolic profiles of hypertensive and control rat pericytes were determined using a Seahorse XFp Extracellular Flux Analyzer (Agilent Technologies, CA, USA). Mitochondrial respiration, glycolysis rate, and real-time ATP production rate were measured using suitable kits (Cell Mito Stress Test, Glycolysis Stress Test, Real –Time ATP Rate Assay) according to the manufacturer’s instructions and as already shown^49^. Briefly, after cell harvesting, pericytes were resuspended in culture medium and seeded at 1.5 x 10^4^ cells/well in Seahorse cell culture microplates and incubated overnight in a CO_2_ incubator. After replacing the growth medium with pre-warmed assay XF DMEM medium pH 7.4 supplemented with 1 mM pyruvate, 2 mM glutamine, and 10 mM glucose for Mito Stress Test and ATP production rate and only with 2 mM glutamine for Glycolysis Stress Test, the cells were preincubated at 37 °C for 45 min. in a non-CO_2_ incubator. The cartridges were loaded with assay medium and with standard inhibitors/substrates: 5 µM oligomycin, 2.25 µM FCCP, and 1 µM rotenone/ antimycin A mixture for Mito Stress Test and with 10 mM glucose, 5 µM oligomycin, and 50 mM 2-deoxy-glucose for Glycolysis Stress Test or with 5 µM oligomycin and 1 µm rotenone/anitmycin A mix for ATP production rate measurement. For experiments with metabolic inhibitors first port of cartridge was loaded with cocktail of mitochondrial inhibitors (2 µM UK5099, 3 µM BPTES, 4 µM Etomoxir or glycolysis inhibitor (50 mM 2-DG) and last 3 ports with standard inhibitors. After the cartridges were calibrated, measurements were initiated. For ATP production rate measurements, the assay medium in cell microplates was again changed to fresh one. The Seahorse assays were analyzed using XF Wave 2.6.1 software, according to manufactureŕs instructions. After the measurements, cells were collected and lysed using RIPA buffer, and diluted with PBS. Total protein concentrations were measured on an absorbance microplate reader Sunrise (TECAN, Switzerland) using BCA Protein Assay (Sigma-Aldrich, CA, USA). For all calculations, oxygen consumption rate (OCR) and extracellular acidification rate (ECAR) were normalized to the total amount of protein in each well and expressed per μg of protein. Cell Mito Stress Test was used to investigate most important mitochondrial properties: proton leak, ATP linked respiration, maximal respiration, and spare respiratory capacity. Glycolysis Stress Test was used to study glycolytic function of pericytes, and ATP test to explore real-time ATP production rate in control and hypertensive pericytes.

### Statistical Analysis

Data analysis was performed using GraphPad Prism 10 software. All data are shown as mean ± standard error of the mean (s.e.m), and exact numbers for each dataset are detailed in figure legends. Data were assessed for normality. Parametric unpaired t-tests were used to compare normally distributed data between two groups. For comparison between multiple groups, a one-way ANOVA was used followed by Holm-Sidak to correct for multiple comparisons. For comparison between multiple groups and conditions, the data were analyzed using two-way ANOVA followed by Holm-Sidak post hoc correction. *P*-values ≤ 0.05 were considered to indicate statistical significance.

## Results

### Stage-specific brain vascular cells adaptation to hypertensive stress

In our cross-sectional study, we examined the effects of hypertension across three defined stages: initial hypertension (HTN), early chronic HTN, and late chronic HTN stages in the SHRSP model, on brain microvascular cells (Fig. 1a). We identified brain vascular cells, particularly focusing on differentiating endothelial cells (CD31^+^) from pericytes (PDGFRß^+^) (Fig. 1b). Our initial identification of pericytes was based on the positive selection of PDGFRß, a marker that is expressed in all pericyte subtypes^40^, and subsequently subdivided pericytes based on their expression of NG2 and CD13^40^. Our results showed that at the onset of HTN, the frequency of CD31^+^ endothelial cells and the vascular pericyte compartment remained consistent between groups (Fig. 1c-e).

**Fig. 1.**
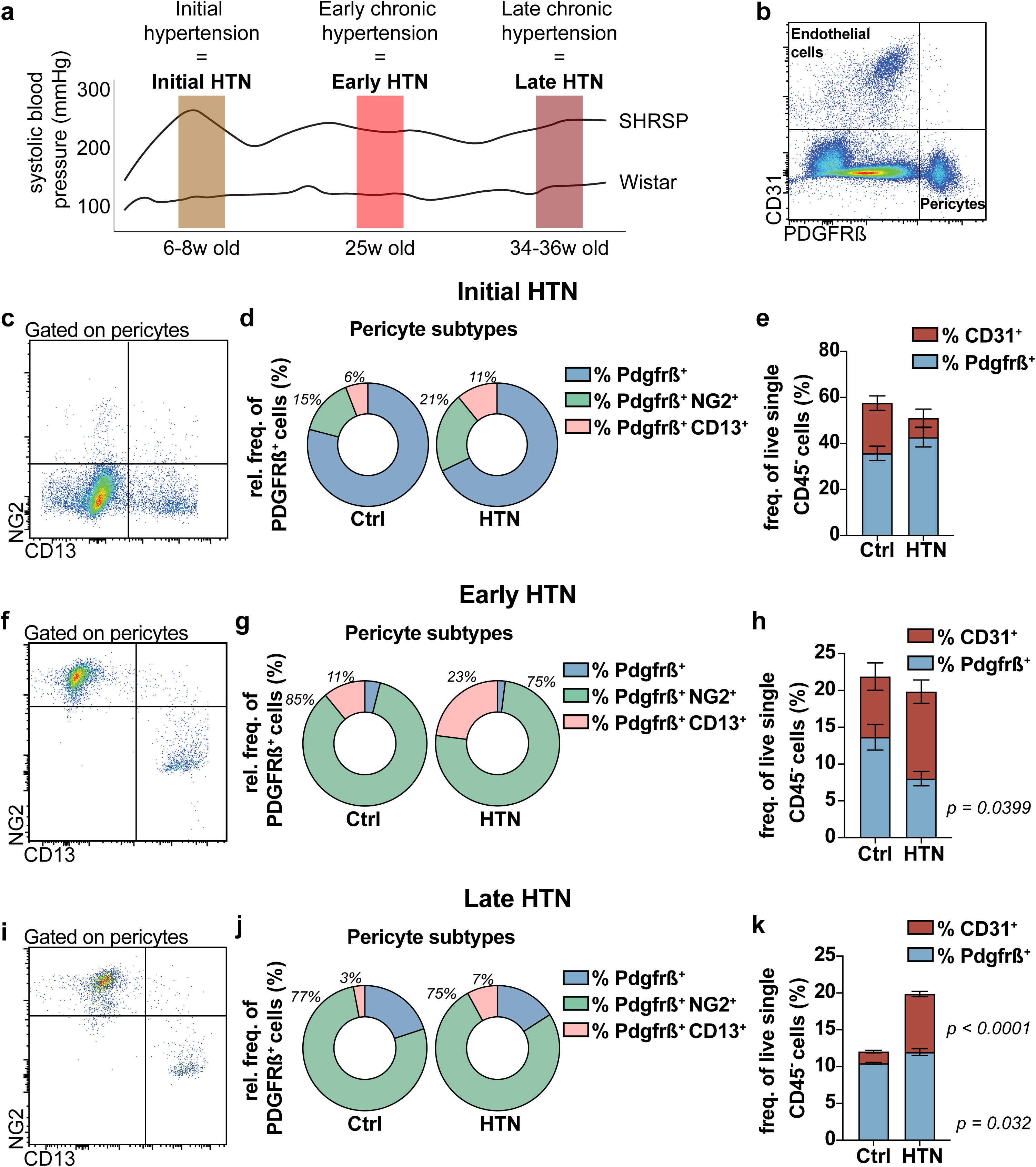
Progressive phenotypic shifts in brain pericytes across arterial hypertensive states. Representative depiction of systolic blood pressure across three hypertension phases in SHRSP and Wistar rats. (a) Systolic blood pressure (mmHg) in SHRSP rats demonstrate increasing hypertension (HTN) from initial HTN (6-8 weeks), early chronic HTN (25 weeks), and late chronic HTN (34-36 weeks) phases, with Wistar rats as controls. Age of rats at each time point is shown on the x-axis. Blood pressure trends are representative, detailed in ^29^. (b) Gating strategy employed to select live vascular cells, excluding CD45 cells, and subsequent identification of CD31^+^ endothelial cells and PDGFRß^+^ pericytes. (c-k) Pericyte subtype identification across hypertension phases: initial HTN (c-e), early chronic HTN (f-h), and late chronic HTN (i-k). Differential expression of NG2 and CD13 among PDGFRß+ pericytes is depicted (d, g, j), alongside relative frequencies and comparisons of endothelial and pericyte populations (e, h, k). For experimental methodology see Suppl. Fig. 1a, b. n=6 per group for initial and early chronic HTN, n=5 per group for late chronic HTN; experiments were performed twice per group per time point. Data are presented as mean ± s.e.m.

During early chronic HTN (Fig.1f, g), the vascular system adapts to persistent hypertensive conditions, resulting in enhanced expression of NG2 and double the amount of CD13 pericytes compared to controls (11% vs. 23%, *p* = 0.0335). A significant decrease was observed in frequency of PDGFRß^+^ cells in HTN (13% vs. 8%, *p* = 0.0399) (Fig. 1h), while the frequency of CD31^+^ endothelial cells remained stable between groups. This result showed a specific susceptibility of pericytes to early chronic HTN stress (Fig. 1h).

In the late chronic HTN phase, NG2 expression dominated the pericyte population (Fig. 1i, j). Quantitative analysis revealed that in controls, only a small fraction of PDGFRß^+^ cells co-expressed CD13 (3%), while a substantial majority co-expressed NG2 (77%). In contrasted to early chronic HTN, CD13 co-expression decreased to 7%, NG2 co-expression remained the same, and PDGFRß^+^ expression increased, marking a nuanced yet significant shift in pericyte subtypes (Fig. 1j). Furthermore, we observed a significant increase in the frequency of CD31^+^ endothelial cells in late chronic HTN (20% vs. 11% in controls, *p* <0.0001) and a modest rise in PDGFRß^+^ HTN pericytes (12% vs. 10% in controls, *p* = 0.032) (Fig.1k).

### Differential proliferative capacity of brain endothelial cells and pericytes in hypertensive states

We conducted further analyses to investigate the proliferative responses of endothelial cells and pericytes across hypertension phases. In the initial HTN (Fig. 2a-d), while the absolute numbers of endothelial cells remained stable, there was a significant increase in Ki67 expression in the hypertensive group (Fig. 2a, b). Conversely, there were no discernible differences in the absolute numbers of PDGFRß^+^ pericytes or their Ki67 expression between Ctrl and HTN displaying an early resilience in response to hypertensive stress (Fig. 2c, d). In early chronic HTN (Fig. 2e-h), endothelial cells maintained stable numbers without the same proliferative increase observed in pericytes. During this phase, there was a significant reduction in the absolute number of pericytes in HTN alongside an increase in Ki67 expression, depicting pericytes under stress, both decreasing in number and increasing in proliferative activity (Fig. 2g, h), pointing to a unique interplay between the cellular responses of each vascular component. In the late stage of chronic HTN (Fig.2 i-l), while endothelial cells displayed a significant increase in both numbers and Ki67 expression in HTN, our results showed a significant increase in the number of pericytes and a decrease in Ki67 expression.

**Fig. 2.**
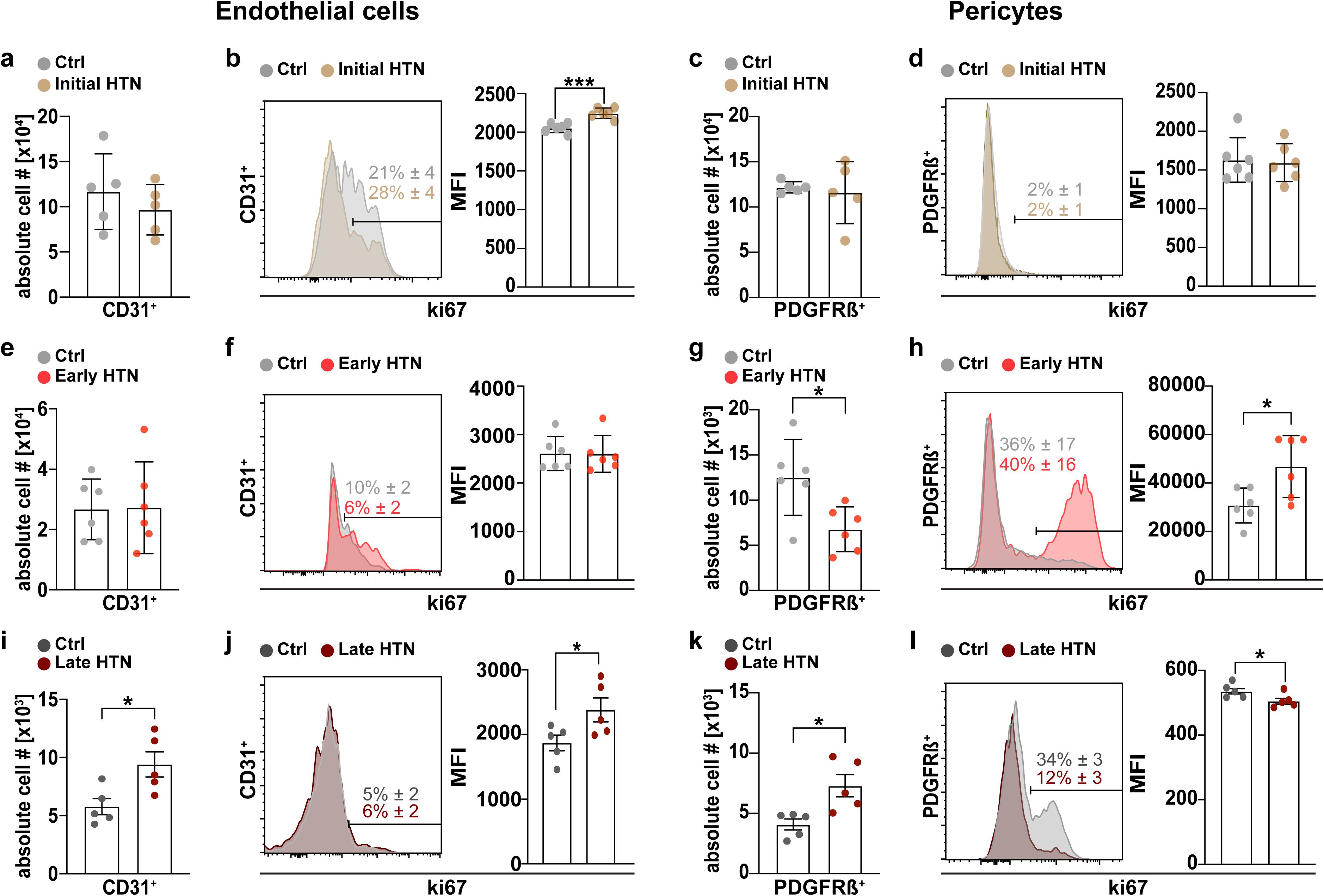
Vascular cell abundance and proliferation across hypertension stages. Analysis of endothelial cells and pericytes abundance across hypertension stages. Row-wise representation of initial HTN (a-d), early chronic HTN (e-h), and late chronic HTN phases (i-l), detailing absolute numbers of PDGFRß^+^ pericytes and CD31^+^ endothelial cells comparing control (Ctrl) vs. hypertensive (HTN) groups. Histograms and bar graphs represent Ki67 expression and quantification of CD31^+^ and PDGFRß^+^ cells. For experimental methodology see Suppl. Fig. 1a. Each data point represents one biological sample, experiments were performed twice per group and time point. Data are presented as mean ± s.e.m. *p < 0.05, ***p < 0.001.

### Tracing the development of vascular markers in hypertension progression

Building on our comprehension of hypertension-associated vascular remodeling, we aimed to delineate the dynamic vascular profile throughout the progression of hypertension. Using UMAP analysis, we projected CD31^+^ and PDGFRß^+^ cells to discern between shifts in vascular cell properties (Fig. 3a). We set to characterize different aspects of vascular cell behavior that underlie the significant changes observed in cellular marker expression and population dynamics across different stages of hypertension. We investigated the vascular identity of hypertension (Fig. 3b), revealing a multi-faceted landscape where CD31, PDGFRß, NG2, and CD13 expressions converge. This analysis revealed an evolving expression pattern of endothelial and pericyte markers, pinpointing the location of PDGFRß clusters and identifying subclusters expressing NG2 and CD13, which point to the intricate interplay within the vascular cell milieu. Normalizing MFI to the initial control expression levels of CD31, PDGFRß, NG2, and CD13 allowed us to trace the marker expression trajectory throughout hypertension development (Fig. 3c). Initial and late chronic hypertension stages exhibit elevated CD31 expression showing an endothelial component reacting to hypertensive conditions. Conversely, PDGFRß expression peaked in initial controls, while NG2 and CD13 expressions dominated the hypertensive groups, especially in the chronic phases. Lastly, we aimed to encapsulate the spatial and temporal shifts in marker expression profiles to provide a clear visual narrative of each individual marker over time. Our results showed the phenotypic shift in critical vascular markers, illustrating the complex development of vascular identity under the influence of hypertension (Fig. 3d).

**Fig. 3.**
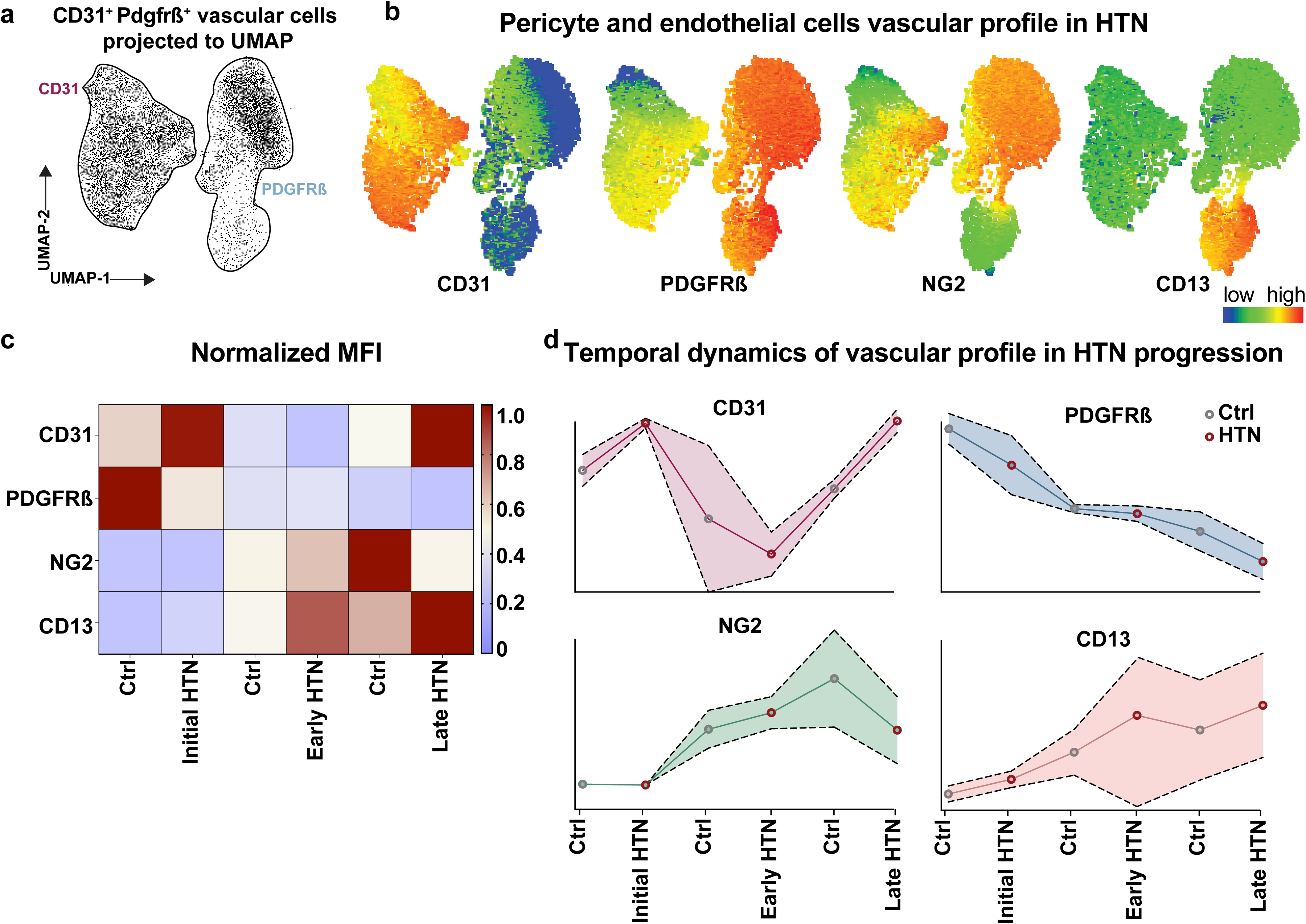
Temporal dynamics of vascular cell marker expression across hypertension progression. UMAP visualization of vascular cell heterogeneity and expression patterns. (a) UMAP projection of CD31^+^ and PDGFRß^+^ cells. (b) Representative expression heatmaps of CD31, PDGFRß, NG2, and CD13 within the vascular clusters to depict overlapping expression of investigated pericyte markers. (c) Heatmap of vascular cells expression levels of CD31, PDGFRß, NG2, and CD13 across six groups: initial control, initial hypertension, early chronic control, early chronic hypertension, late chronic control, and late chronic hypertension, from two independent experiments per time point, shown as normalized mean fluorescence intensity (MFI) values to the expression levels of the initial control group. For experimental methodology see Suppl. Fig. 1a. (d) Line graphs displaying the temporal progression of marker expression for CD31, PDGFRß, NG2, and CD13, individually across all examined time points.

### Molecular pathways underlying hypertension-induced vascular remodeling

To pinpoint the molecular basis of vascular remodeling in hypertension, we conducted an in-depth analysis of isolated microvessels using a custom PCR array focusing on genes within key pathways pivotal to understanding the vascular consequences of chronic hypertensive states—angiogenesis, BBB integrity, hypoxia, inflammation, and specific pericyte markers. This targeted analysis aimed to link transcriptional changes that accompany and possibly precipitate the vascular alterations seen in hypertension. Our results revealed the microvessel transcriptional landscape, pinpointing genes that undergo significant regulation in response to hypertensive stress. In the early chronic phase (Fig. 4a), a notable upregulation of n = 31 genes (fold change > 1.5, *p* < 0.05), including *Agtr1a, Epas1, Timp3, Tek, Notch3, Vegfb*, and *Mmp9*, displayed a state of heightened vascular reactivity and remodeling potentially driven by inflammatory vascular processes displayed by the upregulation of *TNF, Icam1, Ccl2, Ccl5, IL10, Il1b, Ifng, Bdkrb1,* and *Nos2* (Fig. 4a, e, and Suppl. Fig. 03). On the other hand, in late chronic HTN n = 35 genes were downregulated (Fig. 4b, e). This difference displays the specific effects of hypertension on vascular gene expression, demonstrating the progression from an active transcriptional response in the early phase to a more subdued profile in late chronic HTN. Next, we examined the biological pathways that were prominent in the observed transcriptional changes. Our findings revealed that early chronic hypertension is a phase marked by active vascular transformation, with upregulated genes indicating a substantial role of the vasculature in processes including blood flow regulation, lipid response, blood pressure control, angiogenesis, and inflammation (Fig. 4c). The late chronic HTN phase revealed a decrease in the activity of pathways, which may indicate a decline in the ability of blood vessels to adapt to high blood pressure and a potential failure to respond to the hypertensive challenge, characterized by a reduction in the expression of genes involved in lipid metabolism, inflammation, cell proliferation, and programmed cell death (Fig. 4d).

**Fig. 4.**
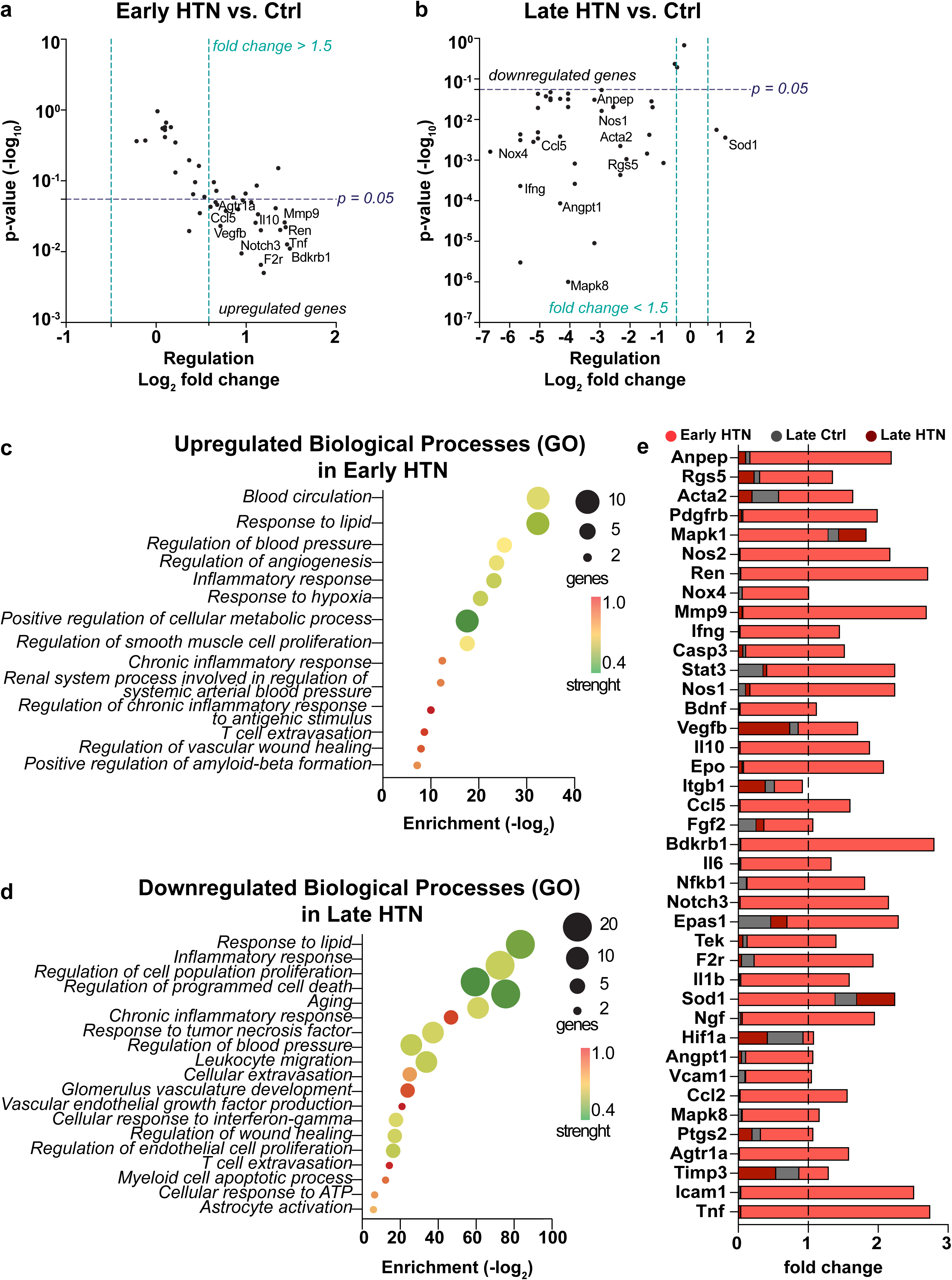
Microvessel-based changes in lipid, angiogenic, and inflammatory pathways during the progression of hypertension. Microvessel isolation methodologies are detailed in Suppl. Fig. 1c. (a) Volcano plot displaying the differential gene expression between control and early chronic hypertensive pericytes. (b) Volcano plot for late chronic hypertensive state showing a predominant downregulation of genes. The vertical green dotted lines indicate the 1.5 fold change threshold. The horizontal blue dotted line indicates the significance threshold (*p* < 0.05). (c-d) Gene Ontology (GO) and reactome pathway analyses displaying the upregulated biological processes in early chronic HTN and downregulated processes in late chronic HTN; exact *p*-values are provided in Suppl. Fig. 04. (e) Side bars representative of individual gene expression levels between early and late chronic hypertensive states against early Ctrl. Relative gene expression levels were normalized to *Hprt* and further normalized to the average expression of early Ctrl.

### *In vitro* pericyte expression dynamics mirror in vivo findings, showing hypertension-driven reductions in PDGFRß alongside elevations in NG2 and CD13

Following the characterization of vascular cell dynamics across chronic hypertensive stages, we shifted our attention to a carefully controlled *in vitro* setting to gain a deeper understanding of the effects of hypertensive stress on pericyte behavior (Fig. 5). Our *in vitro* approach began with the isolation of microvascular fragments, their seeding, expansion, and precise pericyte enrichment, free from endothelial cell influence (Suppl. Fig. 1d, Fig. 2)^44^. Immunofluorescence analyses confirmed the specificity and integrity of our pericyte cultures through PDGFRß expression and paralleled the significant shifts observed in *ex vivo* pericyte marker expression in response to early chronic HTN. *In vitro*, pericytes derived from early chronic HTN displayed a marked upregulation of NG2 and CD13, alongside a reduction in PDGFRß expression. And pericytes derived from early chronic matched controls displayed expression of NG2 and slight expression of CD13, although these control pericytes did not lose PDGFRß expression (Fig. 5a, b). Flow cytometric analysis provided a quantitative perspective on these phenotypic adaptations, underlying the immunofluorescence findings, demonstrating significant changes in the expression pattern of PDGFRß, NG2, and CD13 among pericytes cultured derived from HTN versus Ctrl (Fig. 5c-g). These data reinforces our characterization of the dynamic response of pericytes to hypertensive and environmental stress.

**Fig. 5.**
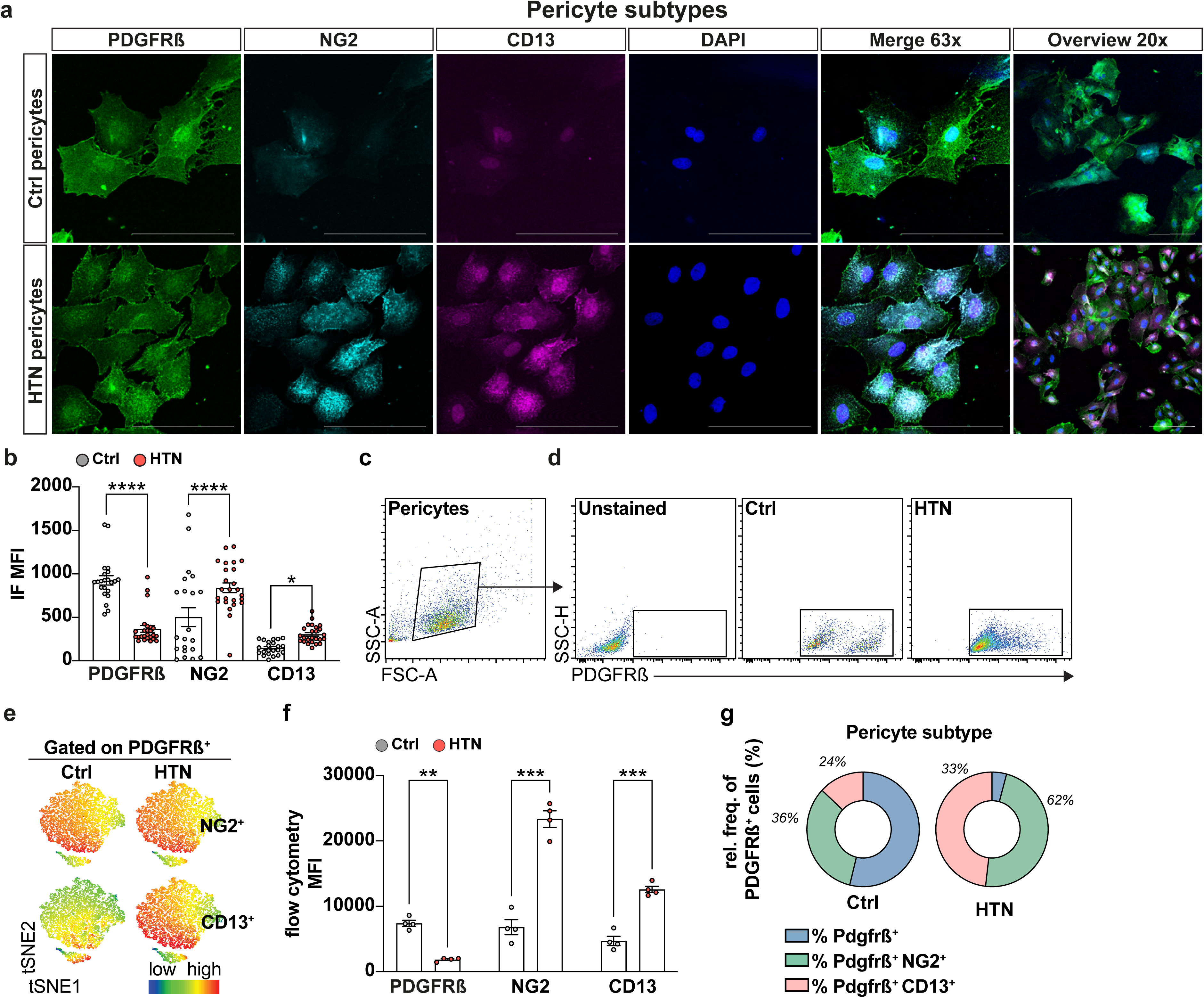
Pericyte expression dynamics revealed a hypertension-driven reduction in PDGFRß and boost in NG2 and CD13 expression. (a) IF imaging analysis for pericyte subtype identification in cultures, revealing differential expression of PDGFRß, NG2, and CD13 across control and hypertensive pericytes. Images show 63x high-resolution merges and a 20x overview of pericyte subtype distribution. (b) Quantitative analysis of mean fluorescence intensity (MFI) for PDGFRß, NG2, and CD13; each dot represents the average MFI measurements obtained per field of view. (c-f) Flow cytometric analysis and tSNE plots elaborating further on pericyte subtype characterization *in vitro*. Initial gating strategies are depicted to validate the expression of PDGFRß in all cells, followed by detailed analysis of NG2 and CD13 expression across pericyte populations using subsequent tSNE visualization and quantification; each dot represents the aggregate data of averaged duplicates derived from *in vitro* expansion that correspond to n=10 biological samples per group. (g) Relative frequencies of PDGFRß^+^ cells displaying subtype distinctions within control and hypertensive groups *in vitro* derived from flow cytometric data. Microvessel isolation and pericyte enrichment methodologies are detailed in Suppl. Fig. 1c, d. Data are presented as mean ± s.e.m. **p < 0.01, ***p < 0.001, ****p < 0.0001. All scale bars represent 100 μm.

### Impact of hypertensive EVs on pericyte mitochondrial function

Expanding on our initial *in vitro* research on pericyte adaptation, we broadened our investigation to examine the impact of hypertensive stimuli on pericyte mitochondrial function at the mitochondrial level, as this is a crucial aspect of maintaining cellular health and vascular integrity during stressful conditions^50^ (Fig. 6). Our methodology centered on isolating plasma and plasma-derived extracellular vesicles (pdEVs) (Suppl. Fig. 1e) from early chronic hypertensive and age-matched control rodents, utilizing these biological materials to simulate normotensive control pericytes *in vitro*. This approach allowed us to directly examine the effects of hypertensive blood components on pericyte mitochondrial function, thereby providing a bridge between systemic hypertension and its vascular repercussions.

**Fig. 6.**
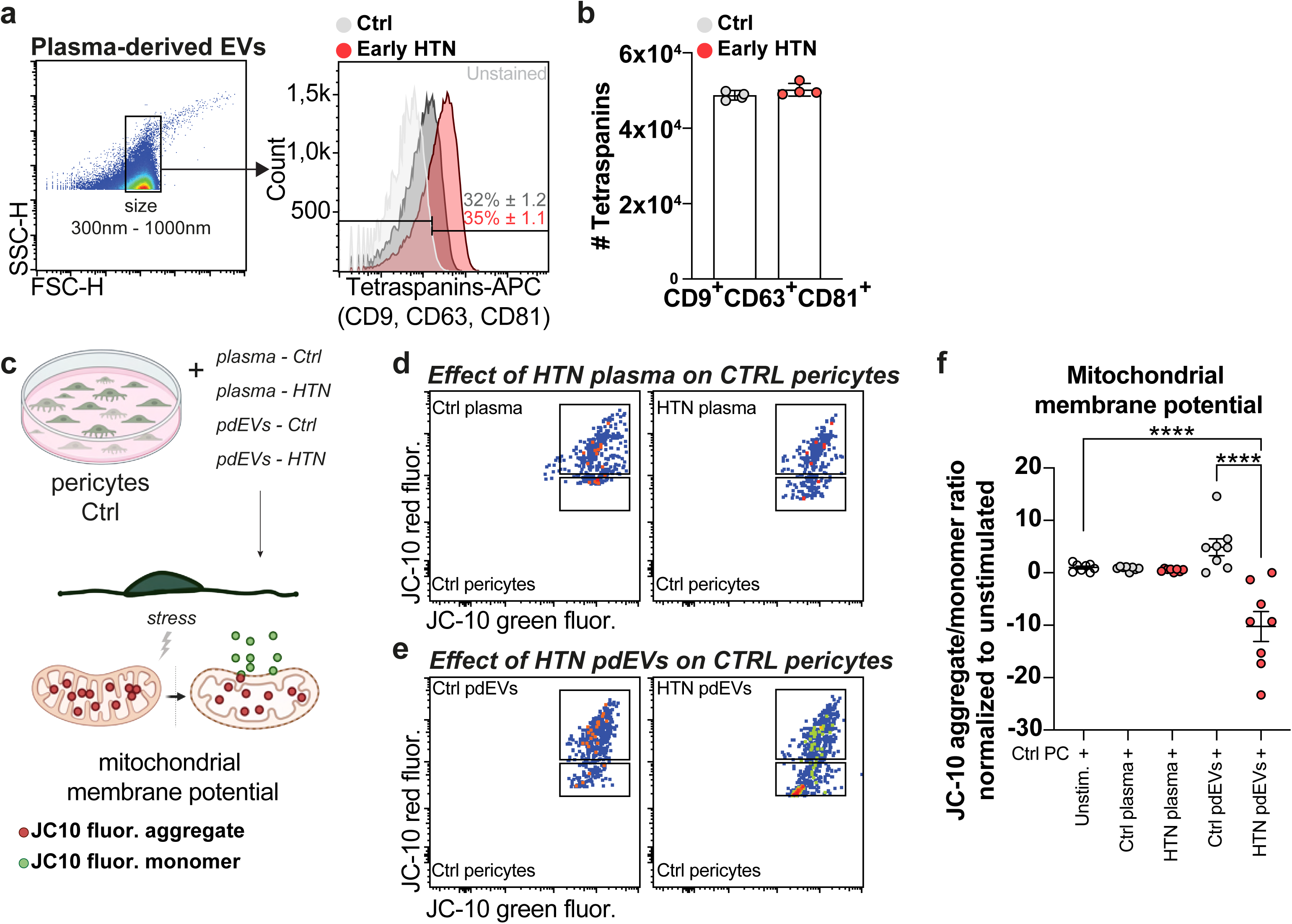
Hypertensive-derived extracellular vesicles cause selective mitochondrial disruption in control pericytes. pdEVs isolation methodologies are detailed in Suppl. Fig. 1e. (a) Flow cytometric analysis displaying EV selection based on size gates using 300-1000nm silica beads, followed by histograms indicating the counts, frequencies, and positive selection of tetraspanins (CD9, CD63, CD81). (b) Bar graphs comparing the absolute numbers of isolated pdEVs from Ctrl and HTN rodents. Each data point represents aggregate data of duplicates derived from one biological sample; experiment was repeated once. (c) Depiction of *in vitro* stimulation conditions with Ctrl pericytes derived from 6w old rodents exposed to plasma or pdEVs derived from early chronic HTN and age-matched Ctrl illustrating the experimental approach using JC-10 staining to assess the impact of hypertensive circulating factors on pericyte mitochondrial function. (d) Flow cytometric plots showing JC-10 fluorescence in Ctrl pericytes stimulated with plasma derived from Ctrl or HTN. (e) Flow cytometric plots showing JC-10 fluorescence in Ctrl pericytes stimulated with pdEVs derived from Ctrl or HTN. (f) Dot plot graph representing the JC-10 aggregate/monomer ratio in Ctrl pericytes under the corresponding stimulation condition normalized to unstimulated pericytes. Each data point represents aggregate data derived from n=10 biological samples measured in duplicates and repeated once. Data are presented as mean ± s.e.m. ****p < 0.0001.

Flow cytometric analysis of pdEVs^48^, with size gating and tetraspanin marker expression selection (CD9, CD63, and CD81), confirmed the presence of isolated pdEVs, revealing no difference in abundance of circulating pdEVs between early chronic HTN and age-matched Ctrl (Fig. 6a, b). However, utilizing the JC-10 dye to assess mitochondrial membrane potential (ΔΨm), a key indicator of mitochondrial health, our experiments revealed a significant impact of hypertensive pdEVs on Ctrl pericyte mitochondrial function (Fig. 6c). Contrary to the relatively stable ΔΨm observed in Ctrl pericytes exposed to normotensive plasma and pdEVs, hypertensive pdEVs induced a marked depolarization of the mitochondrial membrane in control pericytes (Fig. 6d, e). Quantitative analysis of ΔΨm revealed the extent of mitochondrial dysfunction, with hypertensive pdEVs significantly reducing the JC-10 aggregate/monomer ratio, which translates to compromised mitochondria function (Fig. 6f).

### Metabolic reprogramming of pericytes in hypertension-induced vascular cell dysfunction

After uncovering the disruptions to mitochondrial function caused by hypertensive stimuli, we conducted further research on the metabolic characteristics of pericytes derived from hypertensive conditions. We aimed to dissect their broader metabolic response to hypertension-induced stress to capture the essence of their metabolic adaptability and resilience (Fig. 7). Therefore, we conducted an extensive metabolic profile analysis that examined oxidative phosphorylation (OxPhos) and glycolysis in pericytes isolated from initial and chronic stages of hypertension. This approach was predicated on the hypothesis that hypertension has distinct effects on pericyte metabolism throughout its progression, potentially revealing metabolic vulnerabilities or adaptations unique to hypertensive vascular pathology (Fig. 7a-c, Suppl. Fig. 1f).

**Fig. 7.**
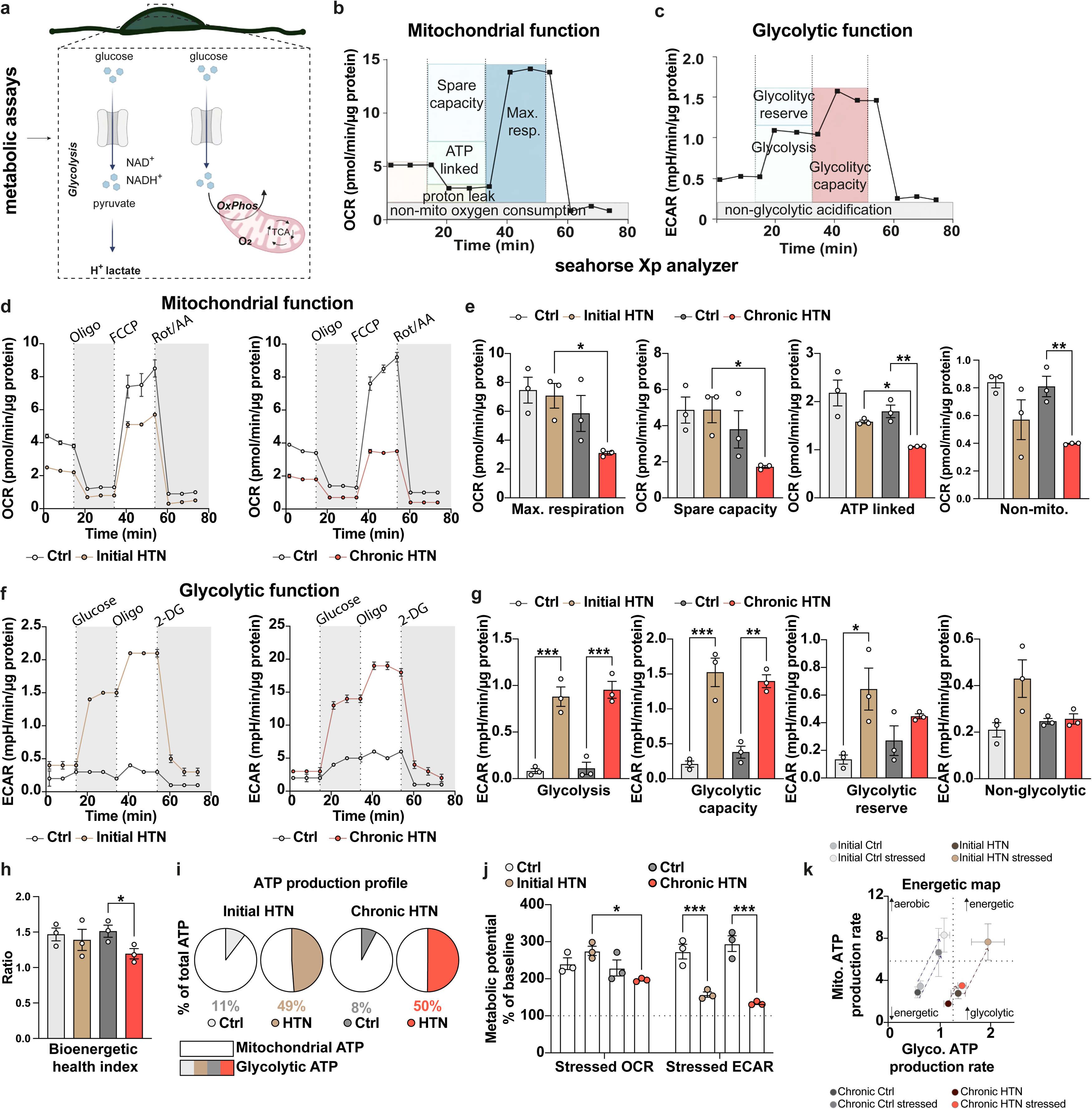
Bioenergetic reprogramming in pericytes upon hypertensive states. (a) Schematic representation of the metabolic pathways analyzed in expanded pericyte cultures from different hypertension phases (Suppl. Fig. 1c, f). (b) Schematic overview of Seahorse extracellular flux analysis measured by mitochondrial stress test in pericytes, highlighting the steps involved in measuring oxygen consumption rate (OCR) and (c) extracellular acidification rate (ECAR). (d) Analyses of OCR in pericytes of initial HTN, chronic HTN, and age-matched controls (*n*= 8-10 per group) using a Seahorse extracellular flux analyzer. Dotted lines indicate the period and time of oligomycin (Oligo), carbonyl cyanide-p-trifluoromethoxyphenylhydrazone (FCCP), rotenone (ROT), and antimycin (Ant) treatment. (e) Bar graphs depicting the maximal respiration capacity, spare capacity, ATP production linked to mitochondrial activity and non-mitochondrial OCR across Ctrl and HTN pericytes. (f) Visualization of glycolytic capacity and reserve in Ctrl and HTN pericytes. Dot lines indicate the period and time of glucose, oligomycin (Oligo), and 2-Deoxy-D –glucose (2-DG) addition. (g) Bar graphs depicting glycolytic capacity, glycolytic reserve, and non-glycolytic ATP production. (h) Bar graph representing the Bioenergetic Health Index as a dynamic measurement of the pericyte response to stress. (i) ATP production profiles in Ctrl and HTN pericytes displaying the relative contribution of glycolysis and mitochondrial respiration to cellular ATP production. (j) Metabolic potential % of baseline showing the metabolic flexibility and potential of Ctrl and HTN pericytes under stress conditions. (k) Energetic map highlighting the metabolic shift in pericytes indicated by the contribution of glycolysis and mitochondrial respiration to ATP production under control and hypertensive conditions. Bar graphs represent mean ± s.e.m. of three independent experiments per group per time point, with each experiment including triplicates. Each data point represent aggregate data from initial groups (Ctrl and HTN) derived from n=8 biological samples each, and chronic groups (Ctrl and HTN) from n=10 biological samples each. Data are presented as mean ± s.e.m. *p < 0.05, **p < 0.01, ***p < 0.001.

Our analyses revealed a pronounced decline in maximal respiration capacity in chronic hypertensive pericytes, apart from their initial hypertensive counterparts (Fig. 7d, e), indicating the progressive impairment of mitochondrial function as hypertension advances. Moreover, evaluation of the spare respiratory capacity, which assesses the potential of cells to respond to bioenergetic stress, showed a significant decrease in chronic HTN (Fig. 7e). The assessment of ATP linked to mitochondrial activity showed a significant decrease in ATP production in pericytes from chronic HTN as compared to both their initial hypertensive counterparts and age-matched controls. This decrease indicates a reduced efficiency of the mitochondria and illustrates a changing energy landscape, where the dependence on ATP synthesis by mitochondria decreases in the presence of hypertensive states (Fig. 7e). Lastly, the notable reduction in non-mitochondrial oxygen consumption rates in chronic hypertensive pericytes showed that the metabolic decline extends beyond mitochondrial function and indicates a general deterioration in cellular metabolic health (Fig. 7e).

Both initial and chronic hypertensive pericytes exhibited significantly higher ATP production via glycolysis, as evidenced by increased ECAR levels (Fig. 7f). This indicates a metabolic shift toward glycolysis in the hypertensive state. The glycolytic capacity (Fig. 7g), reflecting the maximum rate of glycolysis under stressed conditions, was significantly higher in hypertensive pericytes across both stages, emphasizing enhanced glycolytic dependence as a key feature of pericyte metabolic adaptation to hypertension.

Interestingly, the glycolytic reserve (Fig. 7g), which indicates the extent to which cells can further increase glycolysis from their baseline level, showed a significant increase in initial hypertensive pericytes compared to controls. However, this distinction did not persist in chronic hypertension, which represent a potential metabolic limit or saturation in further upregulating glycolysis as the disease progresses. The Bioenergetic Health Index provided a thorough evaluation of the metabolic health of pericytes, revealing a significant decrease in bioenergetic function in chronic hypertensive pericytes. This index emphasizes the negative impact of hypertension on pericyte metabolic capacity (Fig. 7h).

Extending beyond OxPhos and glycolysis measurements, our results revealed a striking shift in ATP production (Fig. 7i). Initial hypertensive pericytes differed significantly from their control counterparts by generating a substantial portion of their ATP through glycolysis, as opposed to control pericytes (49% vs. 11%). This striking shift persisted in chronic HTN (50% vs. 8%). These results show a metabolic shift in hypertensive pericytes, predisposing them to rely more heavily on glycolysis from the onset of the disease. The assessment of metabolic potential under stress conditions validated the influence of hypertension on pericyte energy dynamics (Fig. 7j). Chronic hypertensive pericytes demonstrated a substantially reduced metabolic capacity compared to their initial hypertensive counterparts, implying a limited ability to fulfill increased energy demands or recover from stress. This compromised state was also apparent when contrasting ECAR levels during stress, showing that both initial and chronic hypertensive pericytes were less capable of enhancing glycolysis in response to stress than controls, which revealed a vulnerability in their metabolic response mechanisms. The energetic map illustrates the metabolic states of pericytes across various conditions (Fig. 7k). This visualization emphasizes that initial hypertensive pericytes are primarily glycolytic and exhibit heightened energetic responses when stressed, propelling them into a high-energy production state. In contrast, their control counterparts maintained aerobic profiles under stress conditions.

Given the crucial role of metabolic flexibility in preserving cellular and vascular health under hypertensive stress, we further investigated the consequences of metabolic inhibitors on ATP production in pericytes (Fig. 8, Suppl. Fig. 4). Utilizing mitochondrial inhibitors such as UK5099, etomoxir, and BPTES, we investigated the dependency of pericytes on ATP production pathways derived from mitochondria (Fig. 8a). This resulted in a decrease in spare and maximum respiratory capacity in both Ctrl and HTN pericytes (Suppl. Fig.4a-d), indicating successful mitochondrial pathway inhibition.

**Fig. 8.**
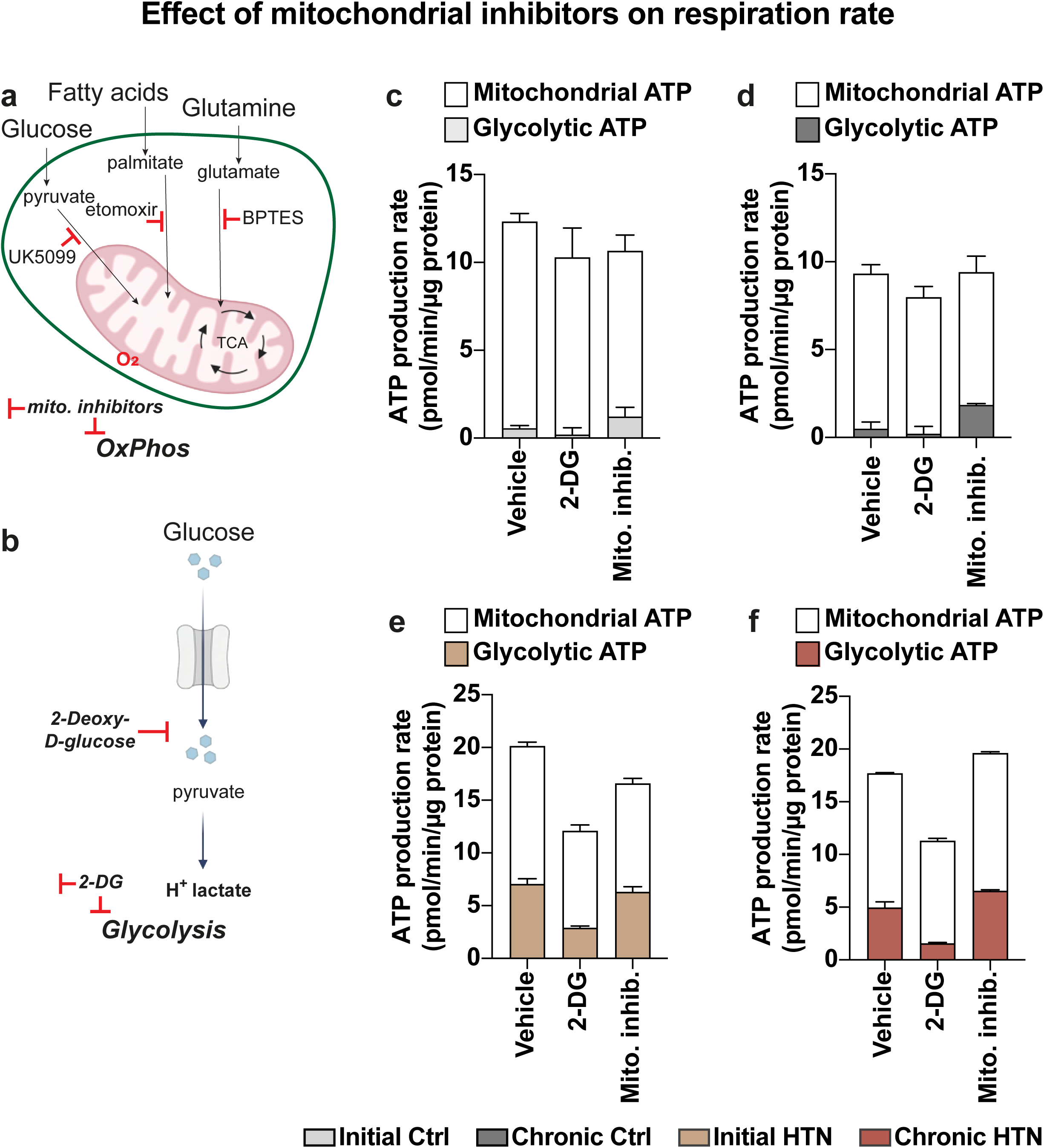
Effect of metabolic inhibition on ATP production in hypertensive and control pericytes. (a) Schematic representation of the application of mitochondrial inhibitors (UK5099, etomoxir, BPTES) to delineate pathways leading to ATP production via mitochondrial oxidative phosphorylation in pericytes. (b) Schematic representation of glycolysis inhibition using 2-deoxy-glucose (2-DG) to block hexokinase. (c, d) Bar graphs representing total ATP production rates measured in Ctrl pericytes showing a predominant dependence on mitochondrial ATP. (e, f) Bar graphs representing total ATP production rate analysis in hypertensive pericytes demonstrating a continued preference for glycolysis. Measurements were performed in triplicate for each group at each time point, with initial groups (Ctrl and HTN) derived from n=8 biological samples each, and chronic groups (Ctrl and HTN) from n=10 each.

In control pericytes, the innate preference for mitochondrial ATP (OxPhos) as the primary ATP production pathway displayed metabolic efficiency and flexibility. In response to mitochondrial inhibitors, these cells effectively increased glycolysis, highlighting their ability to adjust their energy production methods swiftly in response to bioenergetic stressors. Our results showed that control pericytes exhibit an aerobic profile evidence by their reliance on mitochondrial ATP under steady conditions, yet displaying an inherent capacity to engage glycolysis when necessary (Fig. 8c, d, Suppl. Fig. 4e-h).

Considering that hypertensive pericytes rely on glycolysis as their primary energy source, we sought to reduce their primary energy production pathway using 2-DG, a glycolysis inhibitor (Fig. 8b). Hypertensive pericytes exhibited a greater dependency on glycolysis for ATP production, even before being subjected to the same mitochondrial constraints. Inhibition of glycolysis with 2-DG presented a significant bioenergetic challenge, as evidenced by a substantial reduction in total ATP production (Fig. 8e, f, Suppl. Fig. 4k). Although these cells have the potential to shift toward OxPhos, the transition does not fully compensate for glycolytic blockade, highlighting a critical metabolic inflexibility. This observation is crucial, especially considering that initial hypertensive pericytes possess the mitochondrial machinery capable of ATP production, yet they display a marked preference for glycolysis (Suppl. Fig. 4i-l). Upon glycolysis inhibition, chronic HTN pericytes exhibited reduced glycolytic and limited mitochondrial ATP production, displaying an entrenched metabolic programming that favors glycolysis despite the availability of mitochondrial pathways.

## Discussion

In the field of brain vascular biology, our understanding of hypertensive vascular pathology has advanced significantly; however, the phenotypic and metabolic mechanisms of such changes, particularly at the level of cerebral pericytes, remain underexplored. Our *in vivo* and *in vitro* studies have demonstrated significant phenotypic changes in pericytes under hypertensive conditions, characterized by increased co-expression of NG2 and CD13, indicative of a stressed vascular phenotype adapting to ongoing hypertensive insult. These cellular transformations correlate with a profound metabolic reprogramming where pericytes shift toward a glycolysis-dominant metabolism already in the initial phase of hypertension emphasizing a paradigm shift in the metabolic programming of brain pericytes, which, although energy-efficient under hypoxic or stress conditions, delineates a form of metabolic inflexibility when faced with prolonged hypertensive insult. This shift is not merely a passive reaction to altered physiological conditions but a proactive adaptation that can profoundly influence vascular metabolism and function^51, 52^.

Pericytes, traditionally recognized for their role in capillary blood flow regulation and BBB integrity^7,16, 20^, emerged from our study as central players in the metabolic reprogramming associated with hypertension. We detected already in the initial stages of HTN a modest yet significant diversification in pericyte marker expression early on, with increases in NG2 and CD13 co-expression, indicating an early cellular response to rising blood pressure. Our findings revealed a critical transformation in pericyte phenotype, marked by upregulation of NG2 and CD13 and concurrent downregulation of PDGFRß. After central nervous system (CNS) injury, NG2-reactive pericytes can be found near microvessels^13^, where they function as inflammation sensors and assist with immunosurveillance and effector functions of extravasated neutrophils and macrophages. These findings imply that NG2 may play a role in CNS injury that is linked to inflammation. It seems probable that NG2 also plays a role in the development of cSVD, which is characterized by increased BBB permeability, inflammatory infiltrates, and CNS damage^29^. Increased expression of CD13 is a hallmark of an inflammatory response as it facilitates monocyte and endothelial cell adhesion, which is a critical event in the progression of vascular diseases^53^. This finding is consistent with literature showing that CD13 plays a pivotal role in inflammation and vascular remodeling^53, 54^. Specifically, CD13 functions as a mediator of monocyte adhesion to endothelial cells^54^, potentially contributing to the inflammatory environment associated with cSVD. These results indicate a phenotypical pericyte shift from its typical functions toward a phenotype that is prepared for acute stress response and tissue repair. Interestingly, the distinct expression of these markers highlights a specific involvement of NG2 and CD13 in disease-specific vascular remodeling processes. NG2 acts as a general stress response marker, with its increased expression occurring in both normal and hypertensive conditions, whereas the upregulation of CD13 under hypertensive stress emphasizes its role in severe vascular responses, potentially leading to pathological changes.

The gene expression profile of microvessels across various stages of hypertension demonstrates a complex interplay of mechanisms that drive vascular pathology in chronic hypertension. The early stage of chronic hypertension was characterized by an upregulation of genes involved in lipid metabolism, angiogenesis, and inflammatory responses, suggesting an active process of vascular remodeling. Key genes, such as *Agtr1a, Ren,* and *acta2*, enhance systemic blood pressure regulation through their roles in the renin-angiotensin system, directly affecting pericyte contractility and vascular tone. Similarly, *F2r* and *Notch3,* support the restructuring of vascular architecture by influencing both pericyte function and endothelial cell interactions essential for adapting to increased hypertensive stress and maintaining vascular integrity. Parallel to these regulatory genes, a significant vascular inflammatory response was evident from the upregulation of *Tnf, Ccl2, Ccl5, Il1b, Il6, Ifng*, and *Nos2*, genes that orchestrate a robust immune response by mobilizing immune cells and exacerbating vascular inflammation, alongside *Il10*, serving a potential compensatory role to mitigate excessive inflammatory damage, preserving vascular function. Structural remodeling can be further mediated by *Mmp9* and *Vegfb*, with *Mmp9* catalyzing extracellular matrix reorganization essential for vascular repair and *Vegfb* supporting pathological angiogenesis, indicating an adaptive yet potentially maladaptive vascular response.

As hypertension progresses to its late chronic phase, a significant shift occurs with a decline in the expression of many previously upregulated genes, which may suggest a possible exhaustion of metabolic and adaptive capacities. However, the significant upregulation of *Sod1* stands out. Sod1, plays a critical role in mitigating oxidative stress by catalyzing the dismutation of superoxide radicals into oxygen and hydrogen peroxide, and increased expression in the late chronic phase might indicate an elevated oxidative stress environment within vascular cells^55^. This adaptive response could represent a mechanism to counteract increased oxidative damage associated with prolonged hypertension^22^, attempting to preserve cellular function and prevent oxidative injury^56^. Nonetheless, persistent high expression of Sod1 might also reflect the chronic stress state of the vascular cells, potentially contributing further pathological changes if the oxidative burden overwhelms the cell’s antioxidative capacities.

In our exploration of the systemic effects of hypertension on vascular cell biology, we observed profound alterations in mitochondrial function when control pericytes were exposed exclusively to plasma-derived EVs from hypertensive rodents, without the influence of other soluble factors or cytokines. Our results demonstrate that circulating EVs within hypertensive states possess a specific ability to disrupt cellular bioenergetics beyond plasma interactions with primary vascular compartments. The observed decrease in mitochondrial membrane potential in response to hypertensive EVs, but not to plasma alone, indicates the potency and specificity of EVs as carriers of pro-pathogenic signals. This depolarization suggests a direct detrimental effect of hypertensive pdEVs on mitochondrial integrity and function, compromising cellular energy production, indicating for the first time a novel pathway through which hypertension may exert vascular damage.These findings are particularly intriguing, given the established role of EVs in mediating cell-to-cell communication and their potential to carry a multitude of bioactive molecules, including inflammatory mediators, microRNAs, and other non-coding RNAs, which can profoundly influence recipient cell behavior^31, 32^. The ability of hypertensive EVs to impair mitochondrial function in pericytes may reflect a mechanistic pathway through which hypertension exerts systemic effects, leading to disruption of vascular and potentially neurovascular functions. This interaction highlights the broader implications of our findings, linking localized vascular dysfunction to systemic vascular health. In the context of hypertension, where elevated systemic inflammatory markers are common^22^, the impact of circulating EVs becomes a critical factor in the perpetuation of vascular pathology. Specific mitochondrial dysfunction induced by hypertensive EVs could contribute to the metabolic shifts observed in pericytes, favoring glycolysis over oxidative phosphorylation as a survival strategy in response to altered systemic conditions.

We further investigated the metabolic requirements of pericytes and observed that hypertensive pericytes undergo a shift toward a glycolytic phenotype to maintain the increased metabolic demands and mitigate oxidative stress, a strategy reminiscent of cancer cells^57, 58^.

This glycolytic shift in hypertensive pericytes, akin to the Warburg effect observed in cancer cells, indicates that pericytes despite the availability of oxygen, opt for a less efficient pathway to generate ATP for sustaining vital functions under stress ^58, 59^. This shift supports not only cellular survival but also vascular stability and neuronal health, particularly in situations of hypertension. However, unlike tumor cells, which leverage this metabolic shift to support rapid proliferation, pericytes in hypertensive states seem to mainly use glycolysis to sustain their vital functions under stress, thereby supporting vascular stability. This adaptation, while beneficial in the short term for cellular survival, may nevertheless lead to long-term cellular exhaustion and vascular dysfunction. A relevant parallel situation is presented by the Warburg effect^58^, in which, despite the availability of oxygen, hypertensive pericytes increasingly resort to glycolysis, a less efficient pathway for generating ATP, similar to the metabolic preference of cancer cells, to favor rapid proliferation at the expense of metabolic efficiency. The implications of these findings extend beyond mere cellular survival influencing the entire NVU, contributing to the inflammatory processes frequently observed in chronic hypertension. Pericytes, which are responsible for regulating cerebral blood flow, may lose their ability to do so effectively due to altered signaling pathways, including those mediated by nitric oxide and inflammatory mediators such as TNFα and IL-6. As a result, the vascular dysfunction can exacerbate neuronal damage. The chronic activation of this inflammatory cascade disrupts not only pericytes, but also endothelial cells and astrocytes, compounding the deleterious effects on cerebral homeostasis. The glycolytic shift in hypertensive pericytes is particularly evident in mitochondrial dysfunction, which becomes more pronounced in chronic hypertension. Despite their glycolytic bias, chronic hypertensive pericytes face limitations in enhancing their energy production under stress due to mitochondrial inefficiencies. This observation suggests that, at advanced disease stages, hypertensive pericytes operate near their maximal capacity, a critical state that risks compromising cellular integrity under prolonged or severe stress. Our data demonstrate that while early hypertensive pericytes retain some ability to utilize mitochondrial pathways, chronic conditions result in heavy reliance on glycolysis. However, this reliance on glycolysis can contribute to their susceptibility when faced with conditions that limit this energy pathway, indicating a significant shift away from the ability to adapt metabolically. This change in energy strategy points to an adjustment that, while initially compensatory, may become disadvantageous due to reduced mitochondrial function. The metabolic inflexibility of chronic HTN pericytes could be attributed to saturation of mitochondrial capacity, where the demand for ATP and biosynthetic precursors exceeds the ability of mitochondria to function efficiently. This profound metabolic shift in pericytes may influence their interaction with other NVU components such as endothelial cells, neurons, microglia, and astrocytes, potentially exacerbating inflammatory responses and contributing to vascular leakage and BBB dysfunction^29^. The metabolic vulnerability of pericytes demonstrates a key aspect of hypertensive pathology and offers a novel target for therapeutic interventions.

In conclusion, our findings present an evaluation of the physiological and cellular strategies employed by vascular cells in response to chronic arterial hypertension, emphasizing significant phenotypic and metabolic transformations. By understanding and potentially manipulating these broad cellular responses, including metabolic pathways, particularly through targeted interventions that restore metabolic flexibility, some of the severe complications associated with chronic hypertension may be mitigated, and cognitive functions might be preserved. This approach could introduce a new era in the treatment of hypertensive vascular diseases, in which a holistic and metabolic modulation of vascular cells becomes a keystone of therapeutic strategies.

## Concluding remarks

Our study demonstrates that cerebral pericytes in hypertensive states undergo significant phenotypic and metabolic reprogramming, shifting predominantly toward glycolysis as an adaptive response to systemic and localized stress factors. This metabolic shift is notably influenced by plasma-derived EVs from hypertensive rodents, which significantly impair mitochondrial function in control pericytes. The pervasive influence of these EVs emphasize their role not merely as biomarkers but as active participants in the exacerbation of vascular pathology across the NVU. These findings affirm the potential of targeting EVs and metabolic pathways in therapeutic strategies to mitigate the vascular complications associated with chronic hypertension.

## Strengths and Limitations

Our study demonstrated metabolic reprogramming and phenotypic shifts of cerebral pericytes in response to chronic arterial hypertension, however, it is important to recognize certain limitations. The reliance on the SHRSP rat model, while invaluable for mimicking human hypertensive small vessel disease, may not fully capture the complexity of human pathology due to species-specific differences. Furthermore, the exclusive focus on pericytes and extracellular vesicles, although providing detailed insights into their roles in vascular remodeling and mitochondrial dysfunction, limits the scope to these elements without considering the broader cellular interactions within the NVU, which could offer additional mechanisms influencing disease progression. Additionally, while our approach evidences the pivotal role of metabolic shifts towards glycolysis in hypertensive pericytes, the in vitro conditions may not entirely replicate the in vivo microenvironment because pericytes are maintained in ideal conditions, potentially oversimplifying the interaction dynamics between pericytes and other systemic, vascular or neural components. Despite these limitations, our study advances our understanding of hypertensive vascular pathology by indicating how metabolic inflexibility and mitochondrial dysfunction in pericytes contribute to cerebrovascular disease progression. The contribution of our study is significant, offering novel insights into the pathophysiology of cSVD, and pointing to metabolic modulation as a promising therapeutic strategy.

## Acknowledgements

We thank Petra Grüneberg, Dr. Abidat Schneider and Cornelia Garz for their expert technical assistance. Figure 6c, 7a-c, 8a, 8b and Suppl. Fig. 1 were created in ©BioRender – Biorender.com. This study was supported by funds from the DZPG-CIRC.

## Author contributions

L.M., and I.R.D. conceptualized the study; L. M. designed experiments and methodology; L.M., A.P.G., and G.DV. conducted experiments and compiled the data with technical assistance provided by L.E.V, S.H., and P.A.; L.M., A.P.G and G.DV analyzed and interpreted experimental data; L.M. created all figures and wrote the original draft of the manuscript; A.P.G., G.DV., L.E.V., S.H., P.A., S.G.M., S.S and I.R.D. reviewed and edited the manuscript; I.R.D. supervised the study; S. S. and I.R.D. acquired all funding. All authors have read and agreed to the published version of the manuscript.

## Extended Data Files

**Suppl. Fig. 1.**
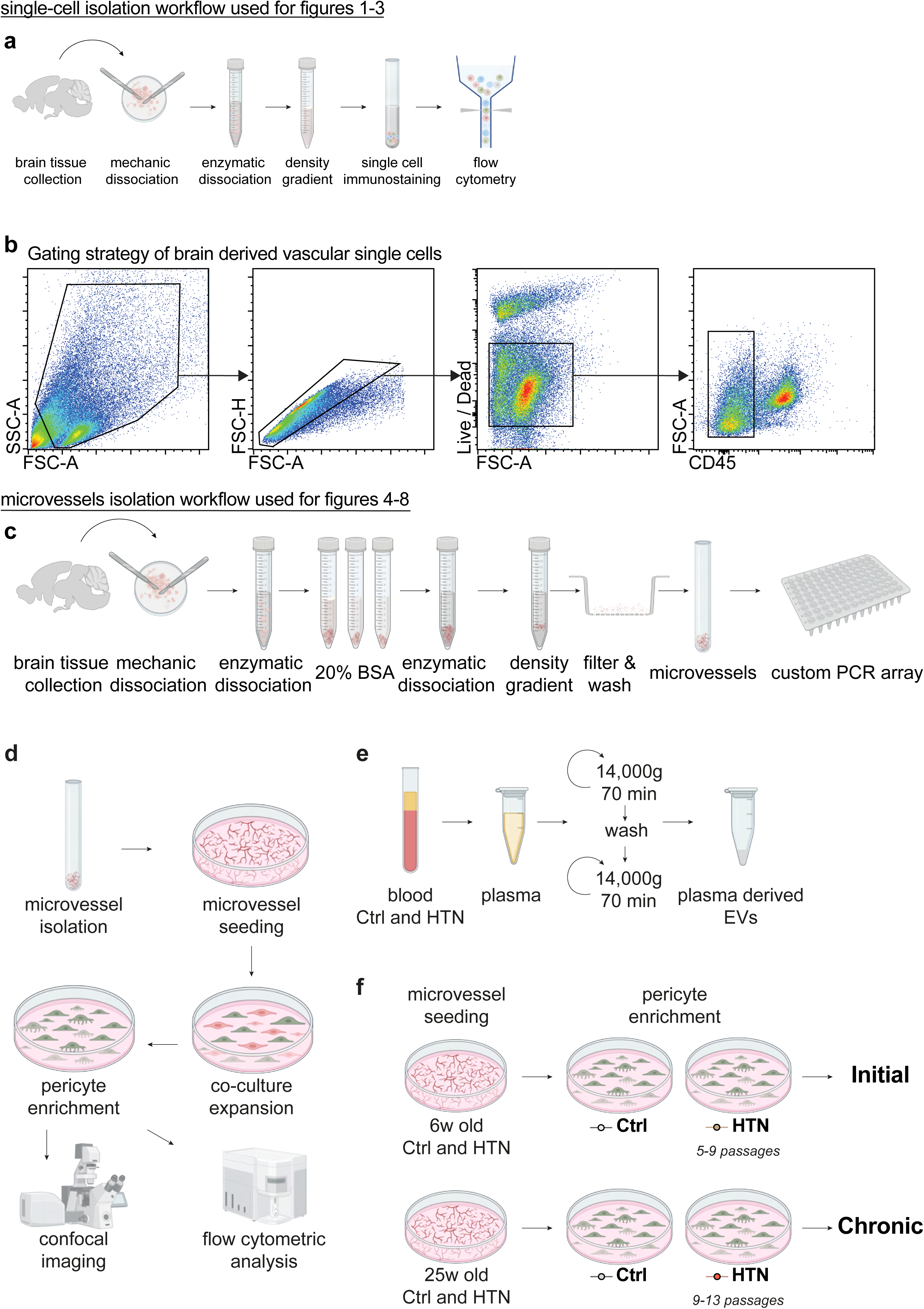
Workflow for isolation and characterization of vascular cells. (a) Summary of single-cell isolation workflow for subsequent flow cytometric analysis. (b) Gating strategy employed to select live vascular cells, excluding CD45^+^ cells. (c) Schematic representation of the multi-step process used for isolating microvascular fragments from hypertensive and control rat brains, followed by RNA extraction for custom PCR array analysis. (d) Schematic representation of the experimental workflow showing initial microvessel seeding, coculture expansion of pericytes with endothelial cells, subsequent pericyte enrichment, and characterization of enriched pericytes through confocal microscopy and flow cytometric analysis. (e) Schematic representation of the experimental setup for isolating and enriching plasma-derived extracellular vesicles (pdEVs) from early chronic HTN and age-matched Ctrl rodents. (f) Experimental design for metabolic pathway analysis of *in vitro* expanded pericyte cultures from different hypertension phases.

**Suppl. Fig. 2.**
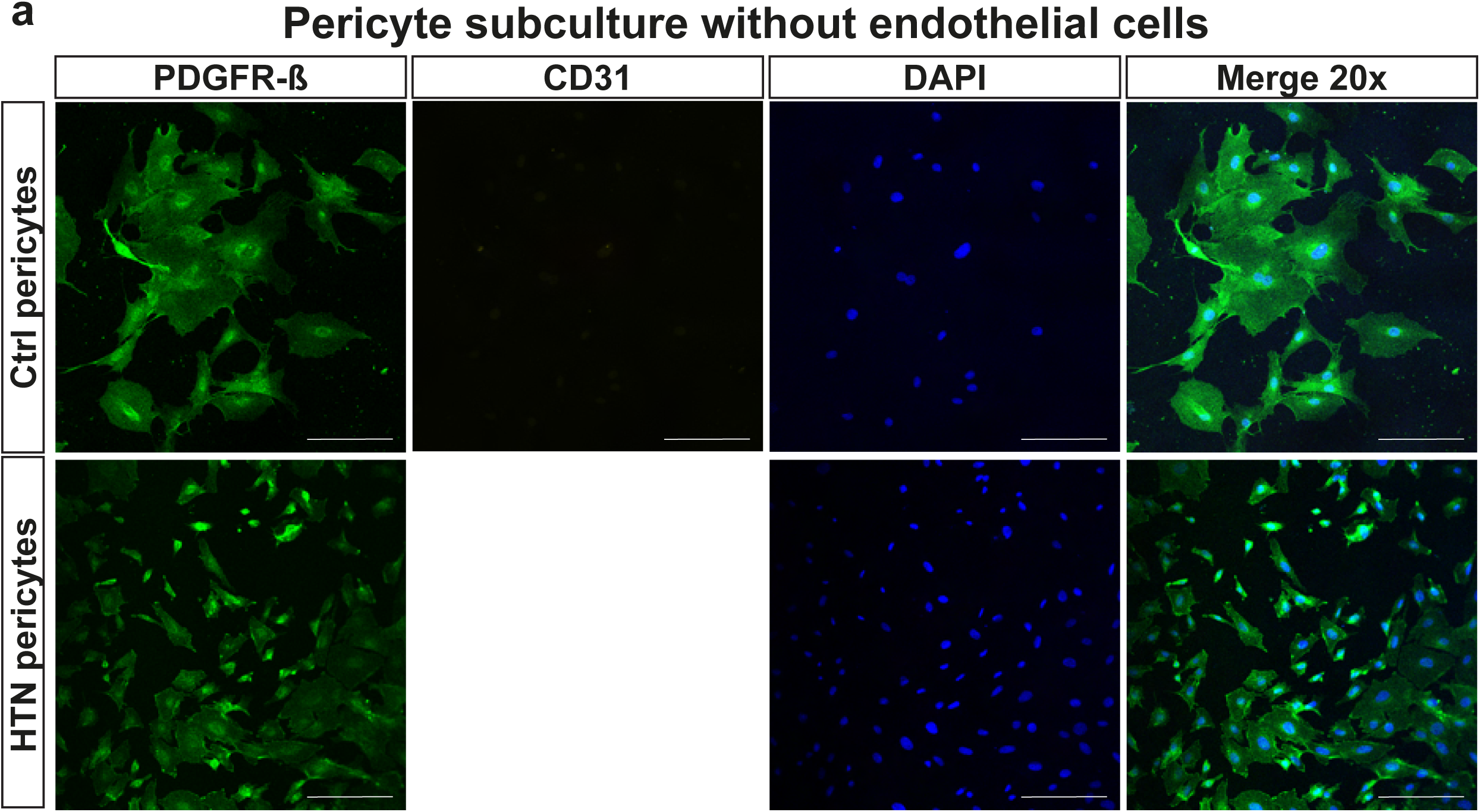
Visualization of pericyte identity and purity by immunofluorescence. (a) Immunofluorescence images displaying pericyte cultures without endothelial cells, with each row depicting control and hypertensive pericytes. Columns depict the expression of PDGFRß, the absence of CD31, and DAPI-stained nuclei, with a 20x merged view. All scale bars represent 100 μm.

**Suppl. Fig. 3.**
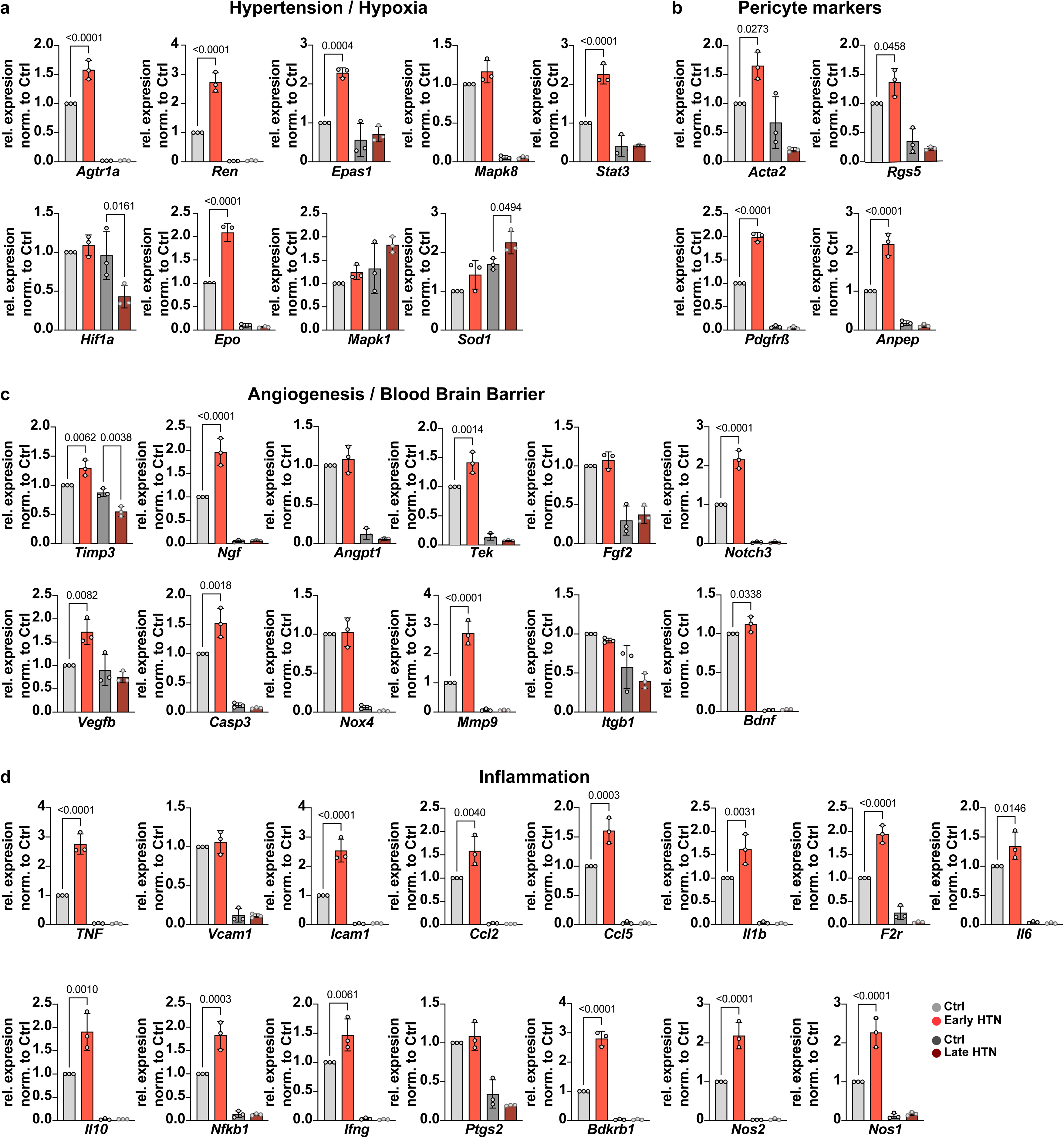
Gene expression profiling in chronic hypertension. Genes within key molecular pathways underlying hypertension-induced vascular remodeling across chronic hypertensive states i.e., (a) hypertension / hypoxia, (b) pericyte markers, (c) angiogenesis / blood-brain barrier, and (d) inflammation. Bar graphs represent mean ± s.e.m. from 3 technical replicates. Each data point represent aggregate data derived from n=5 biological samples.

**Suppl. Fig. 4.**
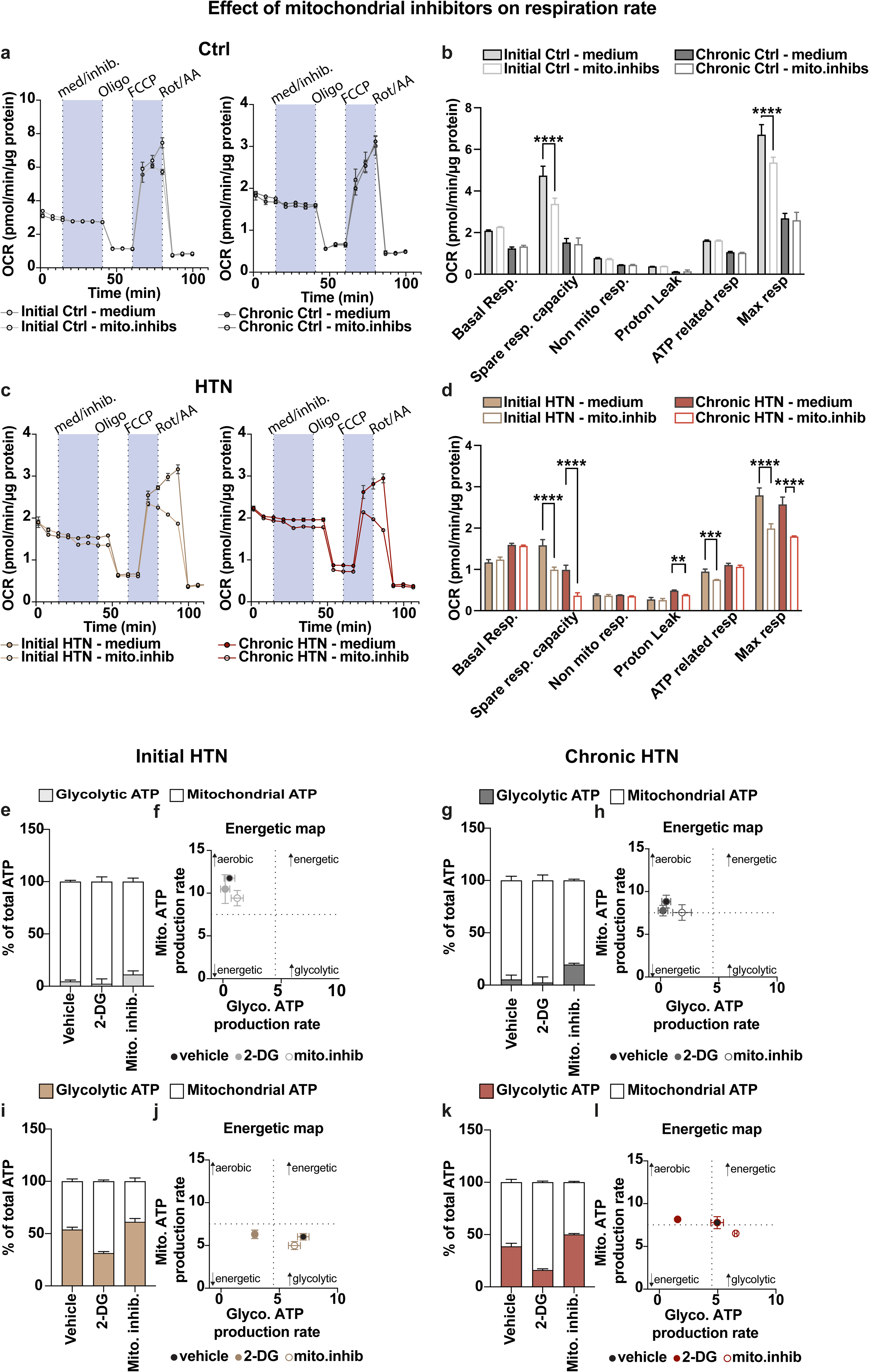
Effect of mitochondrial inhibitors on respiration rate. (a) Oxygen consumption rate (OCR) comparison in initial control pericytes treated with basal medium versus mitochondrial inhibitors. (b) Bar graphs representing the OCR measurements from Seahorse assay in Ctrl pericytes. (c) OCR profiles of HTN pericytes with and without mitochondrial inhibitors across initial and chronic stages of hypertension. (d) Bar graphs representing OCR measurements from Seahorse assay in HTN pericytes. Measurements were performed in triplicate for each group at each time point. Bar graphs represent means ± s.e.m., with initial groups (Ctrl and HTN) derived from n=8 biological samples each, and chronic groups (Ctrl and HTN) from n=10 each. Data are presented as mean ± s.e.m. ****p < 0.0001. (e, g) Bar graphs representing % of total ATP production rates measured in Ctrl pericytes showing a predominant dependence on mitochondrial ATP. (f, h) Energetic maps depicting the aerobic metabolic profiles of Ctrl pericytes across initial and chronic stages. (i, k) Bar graphs representing % of total ATP production rates measured in hypertensive pericytes demonstrating a continued preference for glycolysis. (j, l) Energetic maps depicting the reliance on glycolysis under hypertensive conditions, even when oxidative pathways are available.

**Suppl. Data Table 1.** Experimental overview and sampling distribution across hypertensive phases. Systematic distribution of Wistar and SHRSP rodents used across the various experimental phases and methodologies of the study.

**Suppl. Data Table 01**
| **Experimental Phase** | **Purpose** | **Wistar Rats** | **SHRSP Rats** | **Total per Method** |
| --- | --- | --- | --- | --- |
| Initial Hypertension | FACS Analysis | 6 (2x) | 6 (2x) | 24 |
| Early Chronic Hypertension | FACS Analysis | 6 (2x) | 6 (2x) | 24 |
| Late Chronic Hypertension | FACS Analysis | 5 (2x) | 5 (2x) | 20 |
| Initial Hypertension | In Vitro Culture | 9 | 9 | 18 |
| Early Chronic Hypertension | PCR Array & In Vitro Culture | 10 | 10 | 20 |
| Late Chronic Hypertension | PCR Array | 5 | 5 | 10 |
| Total |  | 58 | 58 | 116 |

**Suppl. Data Table 2.**
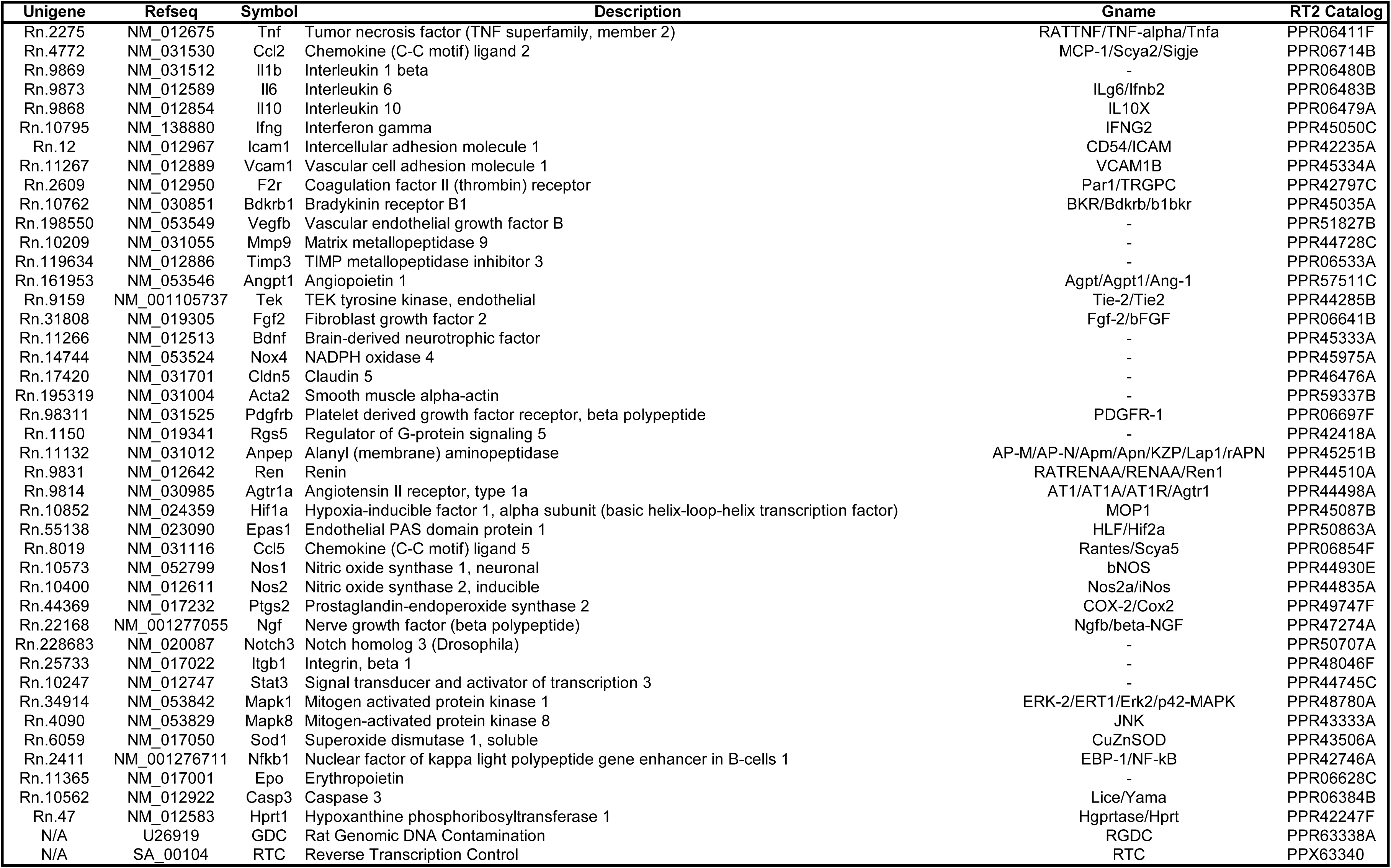
Full gene list of the custom RT^2^ profiler PCR array analysis. List of genes assessed in the custom array profiler (Cat.no. 330171 Custom Array) serving as a resource for our analysis of vascular remodeling in chronic hypertension. The table lists the Unigene identifiers, Refseq numbers, Symbol of each gene, detailed name description, and the RT^2^ catalog numbers specific for each gene.

**Suppl. Data Table 3.** Antibody specifications for vascular cells immunofluorescence analysis. Selection of primary and secondary antibodies employed for the immunofluorescence staining of vascular cells, detailing the target antigen, clone, host species, dilution ratios, and manufacturer details.

**Suppl. Data Table 03**
| **Vascular cell immunofluorescence** |  |  |  |  |  |
| --- | --- | --- | --- | --- | --- |
| Target | Clone | Host | Dilution | Manufacturer | Catalog number |
| CD31 | 390 | Rat | 1:500 | Invitrogen | 14-0311-85 |
| NG2 | D120.43 | Mouse | 1:500 | Invitrogen | 37-2700 |
| PDGFRβ | R.140.4 | Rabbit | 1:500 | Invitrogen | MA5-14851 |
| CD13 | WM15 | Mouse | 1:1000 | BioLegend | 1972330 |
| Anti-Mouse | polyclonal | Goat/IgG | 1:1000 | Invitrogen | A-11001 |
| Anti-Rat | polyclonal | Goat/IgG | 1:1000 | Invitrogen | A-11006 |
| Anti-Rabbit | polyclonal | Goat/IgG | 1:1000 | Invitrogen | A32733 |

## References

1. Yang, A. C. et al. A human brain vascular atlas reveals diverse mediators of Alzheimer’s risk. Nature 603, 885–892 (2022).

2. Obermeier, B., Daneman, R. & Ransohoff, R. M. Development, maintenance and disruption of the blood-brain barrier. Nature Medicine vol. 19 1584–1596 Preprint at 10.1038/nm.3407 (2013).

3. Knox, E. G., Aburto, M. R., Clarke, G., Cryan, J. F. & O’Driscoll, C. M. The blood-brain barrier in aging and neurodegeneration. Molecular Psychiatry vol. 27 2659–2673 Preprint at 10.1038/s41380-022-01511-z (2022).

4. Sweeney, M. D., Zhao, Z., Montagne, A., Nelson, A. R. & Zlokovic, B. V. From Physiology to Disease and Back. Physiol Rev 99, 21–78 (2019).

5. Zlokovic, B. V. The Blood-Brain Barrier in Health and Chronic Neurodegenerative Disorders. Neuron vol. 57 178–201 Preprint at 10.1016/j.neuron.2008.01.003 (2008).

6. Sweeney, M. D., Kisler, K., Montagne, A., Toga, A. W. & Zlokovic, B. V. The role of brain vasculature in neurodegenerative disorders. Nature Neuroscience vol. 21 1318– 1331 Preprint at 10.1038/s41593-018-0234-x (2018).

7. Armulik, A. et al. Pericytes regulate the blood-brain barrier. Nature 468, 557–561 (2010).

8. Daneman, R., Zhou, L., Kebede, A. A. & Barres, B. A. Pericytes are required for bloodĝ€”brain barrier integrity during embryogenesis. Nature 468, 562–566 (2010).

9. Longden, T. A., Zhao, G., Hariharan, A. & Lederer, W. J. Pericytes and the Control of Blood Flow in Brain and Heart. Annual Review of Physiology vol. 85 137–164 Preprint at 10.1146/annurev-physiol-031522-034807 (2023).

10. Hariharan, A., Robertson, C. D., Garcia, D. C. G. & Longden, T. A. Brain capillary pericytes are metabolic sentinels that control blood flow through a KATP channel-dependent energy switch. Cell Rep 41, (2022).

11. Eilken, H. M. et al. Pericytes regulate VEGF-induced endothelial sprouting through VEGFR1. Nat Commun 8, (2017).

12. Török, O. et al. Pericytes regulate vascular immune homeostasis in the CNS. doi:10.1073/pnas.2016587118/-/DCSupplemental.

13. Stark, K. et al. Capillary and arteriolar pericytes attract innate leukocytes exiting through venules and ‘instruct’ them with pattern-recognition and motility programs. Nat Immunol 14, 41–51 (2013).

14. Proebstl, D. et al. Pericytes support neutrophil subendothelial cell crawling and breaching of venular walls in vivo. Journal of Experimental Medicine 209, 1219–1234 (2012).

15. Rustenhoven, J., Jansson, D., Smyth, L. C. & Dragunow, M. Brain Pericytes As Mediators of Neuroinflammation. Trends in Pharmacological Sciences vol. 38 291– 304 Preprint at 10.1016/j.tips.2016.12.001 (2017).

16. Dalkara, T., Gursoy-Ozdemir, Y. & Yemisci, M. Brain microvascular pericytes in health and disease. Acta Neuropathologica vol. 122 1–9 Preprint at 10.1007/s00401-011-0847-6 (2011).

17. Uemura, M. T., Maki, T., Ihara, M., Lee, V. M. Y. & Trojanowski, J. Q. Brain Microvascular Pericytes in Vascular Cognitive Impairment and Dementia. Frontiers in Aging Neuroscience vol. 12 Preprint at 10.3389/fnagi.2020.00080 (2020).

18. Berthiaume, A. A. et al. Pericyte remodeling is deficient in the aged brain and contributes to impaired capillary flow and structure. Nat Commun 13, (2022).

19. Kofler, N. M., Cuervo, H., Uh, M. K., Murtomäki, A. & Kitajewski, J. Combined deficiency of Notch1 and Notch3 causes pericyte dysfunction, models CADASIL, and results in arteriovenous malformations. Sci Rep 5, (2015).

20. Kisler, K. et al. Pericyte degeneration leads to neurovascular uncoupling and limits oxygen supply to brain. Nat Neurosci 20, 406–416 (2017).

21. Sweeney, M. D., Ayyadurai, S. & Zlokovic, B. V. Pericytes of the neurovascular unit: Key functions and signaling pathways. Nature Neuroscience vol. 19 771–783 Preprint at 10.1038/nn.4288 (2016).

22. Ungvari, Z. et al. Hypertension-induced cognitive impairment: from pathophysiology to public health. Nature Reviews Nephrology vol. 17 639–654 Preprint at 10.1038/s41581-021-00430-6 (2021).

23. Fang, C., et al. Arteriolar neuropathology in cerebral microvascular disease. Neuropathology and Applied Neurobiology vol. 49 Preprint at 10.1111/nan.12875 (2023).

24. Schreiber, S., Bueche, C. Z., Garz, C. & Braun, H. Blood brain barrier breakdown as the starting point of cerebral small vessel disease? - New insights from a rat model. Experimental and Translational Stroke Medicine vol. 5 Preprint at 10.1186/2040-7378-5-4 (2013).

25. Evans, L. E. et al. Cardiovascular comorbidities, inflammation, and cerebral small vessel disease. Cardiovascular Research vol. 117 2575–2588 Preprint at 10.1093/cvr/cvab284 (2021).

26. Pantoni, L. Cerebral small vessel disease: from pathogenesis and clinical characteristics to therapeutic challenges. The Lancet Neurology vol. 9 689–701 Preprint at 10.1016/S1474-4422(10)70104-6 (2010).

27. Wardlaw, J. M., Smith, C. & Dichgans, M. Small vessel disease: mechanisms and clinical implications. The Lancet Neurology vol. 18 684–696 Preprint at 10.1016/S1474-4422(19)30079-1 (2019).

28. Quick, S., Moss, J., Rajani, R. M. & Williams, A. A Vessel for Change: Endothelial Dysfunction in Cerebral Small Vessel Disease. Trends in Neurosciences vol. 44 289–305 ePreprint at 10.1016/j.tins.2020.11.003 (2021).

29. Morton, L. et al. Spatio-temporal dynamics of microglia phenotype in human and murine cSVD: impact of acute and chronic hypertensive states. Acta Neuropathol Commun 11, (2023).

30. Blevins, B. L. et al. Brain arteriolosclerosis. Acta Neuropathologica vol. 141 Preprint at 10.1007/s00401-020-02235-6 (2021).

31. Buzas, E. I. The roles of extracellular vesicles in the immune system. Nature Reviews Immunology Preprint at 10.1038/s41577-022-00763-8 (2022).

32. Crewe, C. et al. Extracellular vesicle-based interorgan transport of mitochondria from energetically stressed adipocytes. Cell Metab 33, 1853–1868.e11 (2021).

33. Van Niel, G., D’Angelo, G. & Raposo, G. Shedding light on the cell biology of extracellular vesicles. Nature Reviews Molecular Cell Biology vol. 19 213–228 Preprint at 10.1038/nrm.2017.125 (2018).

34. Bailey, E. L. et al. Differential gene expression in multiple neurological, inflammatory and connective tissue pathways in a spontaneous model of human small vessel stroke. Neuropathol Appl Neurobiol 40, 855–872 (2014).

35. Bailey, E. L. et al. Cerebral small vessel endothelial structural changes predate hypertension in stroke-prone spontaneously hypertensive rats: A blinded, controlled immunohistochemical study of 5- to 21-week-old rats. Neuropathol Appl Neurobiol 37, 711–726 (2011).

36. Held, F. et al. Vascular basement membrane alterations and β-amyloid accumulations in an animal model of cerebral small vessel disease. Clin Sci (2017).

37. Jandke, S. et al. The association between hypertensive arteriopathy and cerebral amyloid angiopathy in spontaneously hypertensive stroke-prone rats. Brain Pathology 28, 844–859 (2018).

38. Kaiser, D. et al. Spontaneous white matter damage, cognitive decline and neuroinflammation in middle-aged hypertensive rats: An animal model of early-stage cerebral small vessel disease. Acta Neuropathol Commun 2, (2014).

39. He, L. et al. Analysis of the brain mural cell transcriptome. Sci Rep 6, (2016).

40. Vanlandewijck, M. et al. A molecular atlas of cell types and zonation in the brain vasculature. Nature 554, 475–480 (2018).

41. Schreiber, S. et al. The pathologic cascade of cerebrovascular lesions in SHRSP: Is erythrocyte accumulation an early phase. Journal of Cerebral Blood Flow and Metabolism 32, 278–290 (2012).

42. Jandke, S. et al. The association between hypertensive arteriopathy and cerebral amyloid angiopathy in spontaneously hypertensive stroke-prone rats. Brain Pathology 28, 844–859 (2018).

43. Crouch, E. E. & Doetsch, F. FACS isolation of endothelial cells and pericytes from mouse brain microregions. Nat Protoc 13, 738–751 (2018).

44. Nakagawa, S. et al. A new blood-brain barrier model using primary rat brain endothelial cells, pericytes and astrocytes. Neurochem Int 54, 253–263 (2009).

45. Lee, Y. K., Uchida, H., Smith, H., Ito, A. & Sanchez, T. The isolation and molecular characterization of cerebral microvessels. Nat Protoc 14, 3059–3081 (2019).

46. Livak, K. J. & Schmittgen, T. D. Analysis of relative gene expression data using real-time quantitative PCR and the 2-ΔΔCT method. Methods 25, 402–408 (2001).

47. Shihan, M. H., Novo, S. G., Le Marchand, S. J., Wang, Y. & Duncan, M. K. A simple method for quantitating confocal fluorescent images. Biochem Biophys Rep 25, (2021).

48. Garza, A. P. et al. Initial and ongoing tobacco smoking elicits vascular damage and distinct inflammatory response linked to neurodegeneration. Brain Behav Immun Health 28, (2023).

49. Debska-Vielhaber, G. et al. Impairment of mitochondrial oxidative phosphorylation in skin fibroblasts of SALS and FALS patients is rescued by in vitro treatment with ROS scavengers. Exp Neurol 339, (2021).

50. Kirkman, D. L., Robinson, A. T., Rossman, M. J., Seals, D. R. & Edwards, D. G. Mitochondrial contributions to vascular endothelial dysfunction, arterial stiffness, and cardiovascular diseases. American Journal of Physiology - Heart and Circulatory Physiology vol. 320 H2080–H2100 Preprint at 10.1152/AJPHEART.00917.2020 (2021).

51. Biswas, S. K. Metabolic Reprogramming of Immune Cells in Cancer Progression. Immunity vol. 43 435–449 Preprint at 10.1016/j.immuni.2015.09.001 (2015).

52. Pålsson-McDermott, E. M. & O’Neill, L. A. J. Targeting immunometabolism as an anti-inflammatory strategy. Cell Research vol. 30 300–314 Preprint at 10.1038/s41422-020-0291-z (2020).

53. Pereira, F. E. et al. CD13 is essential for inflammatory trafficking and infarct healing following permanent coronary artery occlusioninmice. Cardiovasc Res 100, 74–83 (2013).

54. Ghosh, M. et al. Molecular mechanisms regulating CD13-mediated adhesion. Immunology 142, 636–647 (2014).

55. Tsang, C. K. wan, Liu, Y., Thomas, J., Zhang, Y. & Zheng, X. F. S. Superoxide dismutase 1 acts as a nuclear transcription factor to regulate oxidative stress resistance. Nat Commun 5, 3446 (2014).

56. Griendling, K. K. et al. Oxidative Stress and Hypertension. Circulation Research vol. 128 993–1020 Preprint at 10.1161/CIRCRESAHA.121.318063 (2021).

57. Ward, P. S. & Thompson, C. B. Metabolic Reprogramming: A Cancer Hallmark Even Warburg Did Not Anticipate. Cancer Cell vol. 21 297–308 Preprint at 10.1016/j.ccr.2012.02.014 (2012).

58. Wang, Y. & Patti, G. J. The Warburg effect: a signature of mitochondrial overload. Trends in Cell Biology vol. 33 1014–1020 Preprint at 10.1016/j.tcb.2023.03.013 (2023).

59. Brier, M. R. et al. Increased white matter glycolysis in humans with cerebral small vessel disease. Nat Aging 2, 991–999 (2022).

